# VIRGO, a comprehensive non-redundant gene catalog, reveals extensive within community intraspecies diversity in the human vagina

**DOI:** 10.1101/660498

**Authors:** Bing Ma, Michael France, Jonathan Crabtree, Johanna B. Holm, Mike Humphrys, Rebecca Brotman, Jacques Ravel

**Affiliations:** Institute for Genome Sciences, University of Maryland School of Medicine, Baltimore, MD 21201, United States

**Keywords:** vaginal microbiome, metagenome and metatranscriptome reference database, non-redundant gene catalog, intraspecies diversity, gene-centric design, protein family catalog, multi-omics data integration

## Abstract

**Background:** Analysis of metagenomic and metatranscriptomic data is complicated and typically requires extensive computational resources. Leveraging a curated reference database of genes encoded by members of the target microbiome can make these analyses more tractable. Unfortunately, there is no such reference database available for the vaginal microbiome.

**Results:** In this study, we assembled a comprehensive human vaginal non-redundant gene catalog (VIRGO) from 264 vaginal metagenomes and 416 genomes of urogenital bacterial isolates. VIRGO includes 0.95 million non-redundant genes compiled from a total of 5.5 million genes belonging to 318 unique bacterial species. We show that VIRGO covers more than 95% of the vaginal bacterial gene content in metagenomes from North American, African, and Chinese women. The gene catalog was extensively functionally annotated from 17 diverse protein databases, and importantly taxonomy was assigned through *in silico* binning of genes derived from metagenomic assemblies. To further enable focused analyses of individual genes and proteins, we also clustered the non-redundant genes into vaginal orthologous groups (VOG). The gene-centric design of VIRGO and VOG provides an easily accessible tool to comprehensively characterize the structure and function of vaginal metagenome and metatranscriptome datasets. To highlight the utility of VIRGO, we analyzed 1,507 additional vaginal metagenomes, uncovering an as of yet undetected high degree of intraspecies diversity within and across vaginal microbiota.

**Conclusions:** VIRGO offers a convenient reference database and toolkit that will facilitate a more in-depth understanding of the role of vaginal microorganisms in women’s health and reproductive outcomes.

## Background

The microbial communities that inhabit the human body play critical roles in the maintenance of health, and dysfunction of these communities is often associated with disease [1]. Taxonomic profiling of the human microbiome via 16S rRNA gene amplicon sequencing has provided critical insight into the potential role of the microbiota in a wide array of common diseases [2–4]. Yet these data routinely fall short of describing the etiology of such microbiome-associated diseases, such as bacterial vaginosis [5, 6], Crohn’s disease [7, 8] or psoriasis [9], among others. This is perhaps because while 16S rRNA gene sequencing can provide species-level taxonomic profiles of a microbial community, it does not describe the genes or metabolic functions that are encoded in the constituents’ genomes. This is an important distinction because strains of a bacterial species have been documented to exhibit substantial diversity in gene content [10], such that their genomes harbor sets of accessory genes whose presence is variable [11, 12]. It is therefore difficult, if not impossible, to infer the complete function of a microbial species in a specific environment using only the sequence of their 16S rRNA gene. As a consequence, to investigate the role of the human microbiome in health and diseases, particular emphasis should be placed on describing the gene content and gene expression of these microbial communities.

Metagenomic and metatranscriptomic profiling are emerging approaches aimed at characterizing the gene content and expression of microbial communities. Results have led to increased appreciation for the important role microbial communities play in human health and diseases [13, 14]. Despite the rapid development and increased throughput of sequencing technologies, current knowledge of the genetic and functional diversity of microbial community is still highly limited. This is due, at least in part, to a lack of resources necessary for the analysis of these massive short read datasets [13, 15]. *De novo* assembly of metagenomic or metatranscriptomic datasets typically requires rather substantial computational resources and complicates integration of metagenomic and metatranscriptomic data.

Accurate, high-resolution mapping of metagenomic or metatranscriptomic data against a comprehensive and curated gene database is an alternative analytical strategy that is less computationally demanding, prone to fewer errors, and provides a standard point of reference for comparison of these data. Development of such curated databases is crucial to further our understanding of the structure and function of microbial communities [15, 16]. In the last two decades, international initiatives such as MetaHit, the NIH funded Human Microbiome Project (HMP) and the International Human Microbiome Consortium (IHMC) were established to generate the resources necessary to enable investigations of the human microbiome, including large reference taxonomic surveys and metagenomic datasets [13, 17]. While multiple 16S rRNA gene catalogs such as RDP [18], SILVA [19], Greengenes [20], and EZBioCLoud [21] exist, there are relatively few curated resources for referencing metagenomes and metatranscriptomes. Those that do exist focus only on the gut microbiome of either humans [16, 22] or animal model species [23, 24]. A definite unmet demand exists for reference gene catalogs for other body sites such as the oral cavity, the skin, and the vagina [25].

In this study, we constructed the human vaginal non-redundant gene catalog (VIRGO), an integrated and comprehensive resource to establish taxonomic and functional profiling of vaginal microbiomes from metagenomic and metatranscriptomic datasets. VIRGO was constructed using 211 *in-house* metagenomes and 53 metagenomes that were generated under the HMP project [26]. The metagenomic data was supplemented with 321 complete or draft genome sequences of urogenital bacterial isolates. The genes identified in the metagenomes and whole genome sequences were further clustered into Vaginal Orthologous Groups (VOGs), a catalog of functional protein families common to vaginal microbiomes. We meticulously curated the gene catalog with taxonomic assignments as well as functional features using 17 diverse protein databases. Importantly, we show that VIRGO provides >95% coverage of the human vaginal microbiome, and it is applicable to populations from North America, Africa and Asia. Together, VIRGO and VOG represent a comprehensive reference repository and a convenient cataloging tool for fast and accurate characterization of vaginal metagenomes and metatranscriptomes. The gene catalog is a compilation of vaginal bacterial species pan-genomes, creating a vaginal “meta-pan-genome”. We further used VIRGO to characterize the amount of intraspecies diversity present in individual vaginal communities. Previous characterization of these communities using either 16S rRNA gene taxonomic profiling or assembly based metagenomic analyses has failed to resolve this diversity. Here we show that vaginal communities contain far more intraspecies diversity than originally expected. This observation challenges the notion that the vaginal microbiota dominated is by one species of *Lactobacillus*, comprised of a single strain, and could have major implications for the ecology of these otherwise low-diversity bacterial communities. Ultimately, VIRGO and its associated analytical framework will facilitate and standardize the analysis and interpretation of large metagenomic and metatranscriptomic datasets thus expanding our understanding of the role of vaginal microbial communities in health and disease.

## Results

### VIRGO is sourced from a comprehensive collection of vaginal metagenomes and bacterial genomes

VIRGO was constructed using sequence data from fully de-identified vaginal metagenomes (n=264) as well as complete and draft genomes of urogenital bacterial isolates (n=321, de-replicated from 416 genomes). The majority (n=211) of the included metagenomes were sequenced in-house from de-identified vaginal swab specimens. Of the ∼18 billion reads generated for these metagenomes, 14.4 billion (79.7%) were identified as human sequences and removed. Interestingly, the proportion of human reads in the vaginal metagenomes was found to vary with community composition. Vaginal metagenomes dominated by *Lactobacillus* spp. had significantly higher proportions of human sequence reads than those from *Lactobacillus* deficient metagenomes (88.7% vs 73.3%; t=-6.6, P < 0.001; **Additional file 1: Figure S1**). Further pre-processing steps culled sequence reads matching rRNA genes and low sequence quality reads, removing another 1.4% reads. Each metagenome was then *de novo* assembled totaling 1.2 million contigs of length > 500bp with a combined length of 2.8 billion bp and an N50 of 6.2 kbp. Additional metagenomic data (n=53) were obtained from the HMP [13, 14] and contributed 40,000 contigs with length > 500bp, comprising 100 million bp of assembled sequence. The *in-house* metagenomes provided 19.5 times more assembled length than the HMP vaginal metagenomes. In addition to the vaginal metagenomes, we also included 321 complete or draft genome sequences of urogenital bacterial isolates, including 139 from HMP and 277 from GenBank and IMG/M (Integrated Microbial Genomes & Microbiomes) [27]. A summary of the metagenomic reads, assembled contigs and genomes included in the construction of VIRGO can be found in **Additional file 2: Table S1**).

Taxonomic analysis of the 264 metagenomes included in VIRGO, revealed that these communities contained 312 bacterial species present in ≥ 0.01% relative abundance (**Additional file: Table S2**). All major vaginal *Lactobacillus* species (*L. crispatus, L. gasseri, L. iners*, and *L. jensenii*), as well as common facultative and strict anaerobic vaginal species such as *G. vaginalis, A. vaginae, P. amnii, P. timonensis, Megasphaera* genomosp*., Mobiluncus mulieris, Mageebacillus indolicus* (aka. BVAB3)*, Veillonella parvula*, among others were identified in the metagenomes. Even BV-associated bacteria that are often only present at low abundance [28] were represented in the metagenomes, including *Finegoldia magna, Peptoniphilus harei, Peptostreptococcus anaerobius, Mobiluncus curtisii, Peptoniphilus lacrimalis, Anaerococcus tetradius, Eggerthella* spp*., Ureaplasma urealyticum, Veillonella atypica, Corynebacterium glucuronolyticum*, among others. The taxonomic profiles of these communities were further shown to encompass the five previously reported vaginal community state types (CSTs) [29], CST I, II, III, IV, and V with frequencies in this set of metagenomes of 18.9%, 3.8%, 20.5%, 48.5%, and 8.3%, respectively (**Additional file 1: Figure S2. Additional File 2: Table S2**). These results highlight the taxonomic breadth of the vaginal bacterial communities included in the construction of VIRGO (**Additional file 1: Figure S3).**

The dataset used to build VIRGO was compiled from vaginal metagenomes that were obtained from North American women. To determine the comprehensiveness of VIRGO, we mapped reads from 91 vaginal metagenomes that were not included in its construction. These metagenomes were obtained from North American, African [30], and Chinese [31] women, allowing us to determine the utility of VIRGO to analyze metagenomes from other populations. Reads from these metagenomes were mapped to the complete and subsets of the sequence contigs used to build VIRGO. More than 99% of the reads from North American metagenomes were able to be mapped to the complete VIRGO dataset, while only ∼55% of these reads mapped to contigs from the HMP vaginal metagenomes subset (**Fig. 1, Additional file 2: Table S3**). This result indicates a lack of genetic diversity in the HMP vaginal metagenomes, which were derived from highly selected and healthy women [32]. Further, despite originating from populations not used in the construction of VIRGO, 96% and 88% of the reads from African and Chinese women mapped to the complete VIRGO dataset. For these two cohorts, 71.7% and 99.9% of the reads that failed to map to VIRGO, also did not have a match in GenBank (**Additional file 1: Figure S4**). These results illustrate the comprehensiveness of VIRGO and its broad application to different populations and ethnicities. It further shows that the bacterial genetic diversity in the vaginal microbiome across populations is somewhat homogenous.

**Figure 1.**
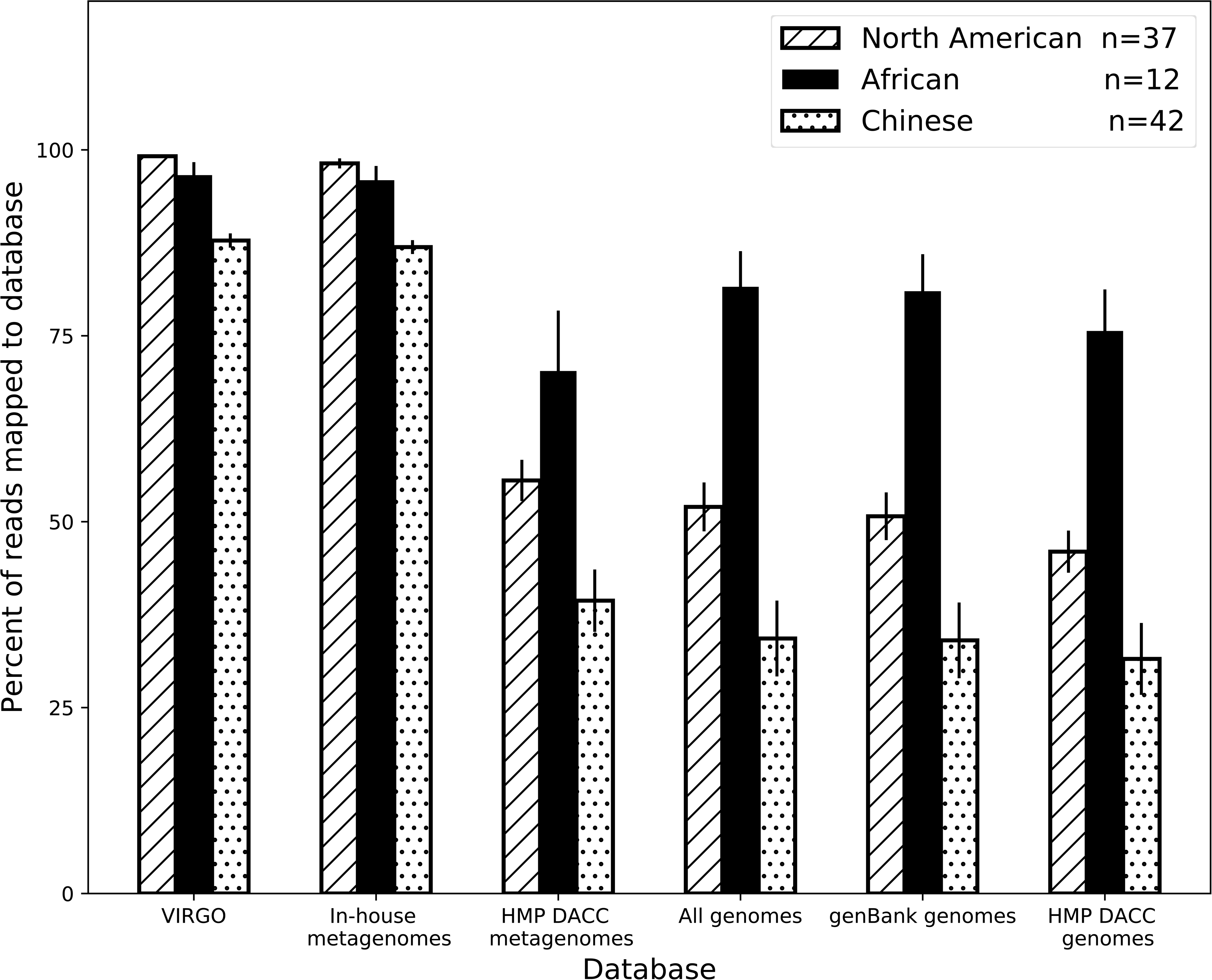
Percent of vaginal metagenome reads that can be mapped to contigs from the following reference data sets: i) complete VIRGO database, ii) 211 *in-house* sequenced vaginal metagenomes, iii) 53 HMP DACC vaginal metagenomes [32], iv) all HMP urogenital reference genomes, v) 277 genomes of bacteria isolated from vagina, reproductive or urinary system deposited in GenBank, and vi) 139 genomes of urogenital bacteria from HMP DACC database [15]. Values plotted are the average, error bars represent the standard error of the mean.

### VIRGO: a non-redundant vaginal bacterial gene catalog

Coding sequences (CDS, n=5,509,298) were predicted from the metagenomic assemblies and genome sequences using MetageneMark [33]. The core workflow to identify and cluster these CDSs is shown in **Fig. 2**, and a more detailed illustration is provided in **Additional file 1: Figure S5.** Metagenomic assemblies contributed ∼80% of the CDSs while the remaining ∼20% of CDSs originated from the urogenital bacteria isolate genome sequences. Redundant genes were then identified and removed via a greedy pairwise comparison at the nucleotide level using highly stringent criteria of 95% identity over 90% of the shorter gene length [16, 22]. This process afforded the removal of partial genes and eliminated overcalling genes as unique because of sequencing errors. A total of 948,158 non-redundant CDSs longer than 99 bp were identified and retained, representing 17.2% of the original 5.5 million CDSs. The *in-house* vaginal metagenomes used to build VIRGO contributed 12 times more non-redundant genes (634,288 genes) than the HMP vaginal metagenomes (54,500 genes). Combined, the metagenomes contributed twice as many non-redundant genes as urogenital bacterial isolate genome sequences (371,099 genes). Metagenomes were found to contain a higher proportion of redundant genes than bacterial genome sequences (84.5% versus 58.1% of their sequence lengths) (**Additional file 2: Table S3**).

**Figure 2.**
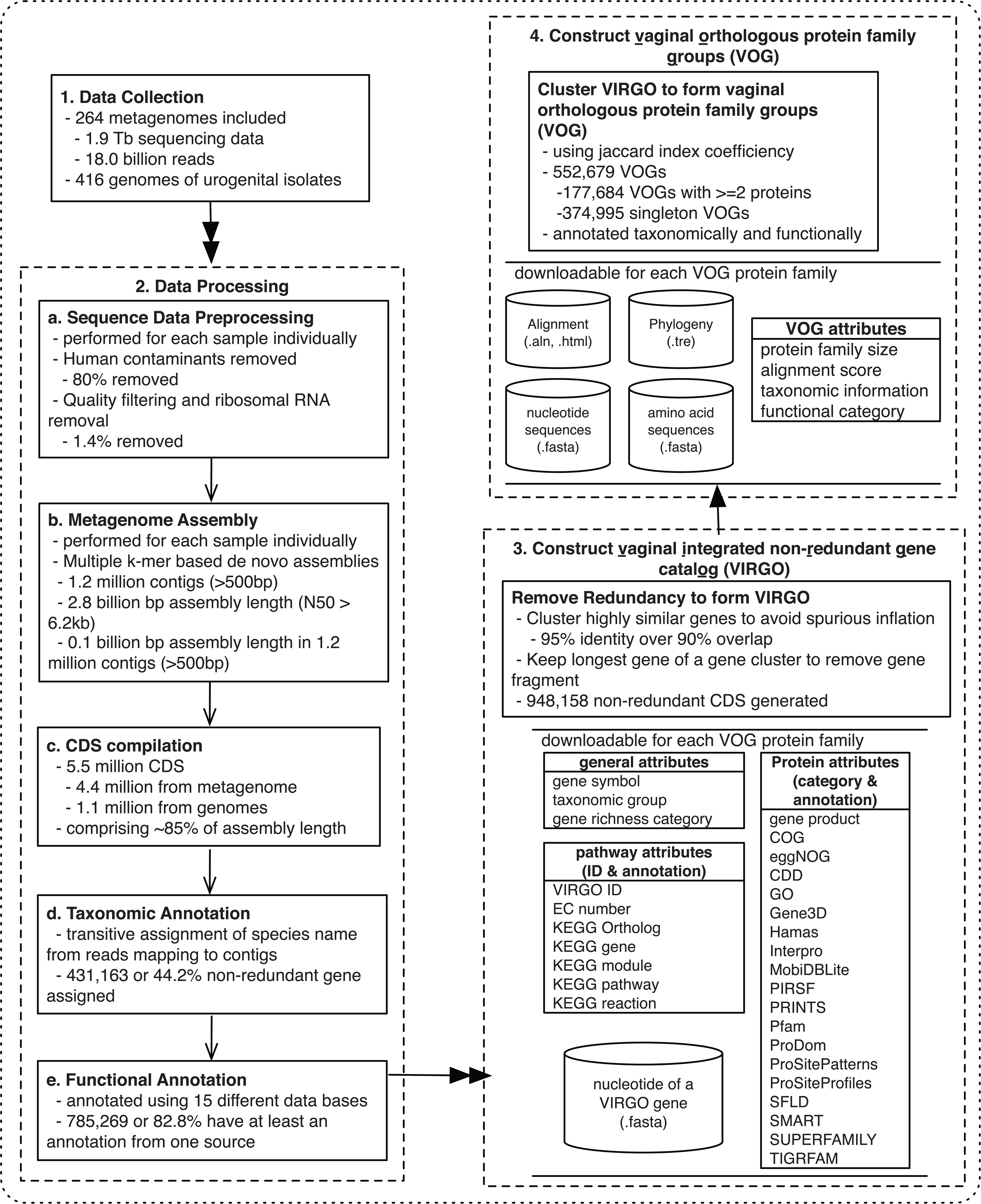
Pipeline for data processing and integration for the construction of the human <u>v</u>aginal integrated non-redundant <u>g</u>ene catal<u>o</u>g (VIRGO) and <u>v</u>aginal orthologous groups (VOG) for protein families. Metagenomes from 264 vaginal metagenomes and 416 genomes of urogenital isolates were processed, that including 211 *in-house* sequenced vaginal metagenomes. The procedures include preprocessing to remove human contaminates, quality assessment, metagenome assembly, gene calling, functional and taxonomic annotation, gene clustering based on nucleotide sequencing similarity to form VIRGO, and jaccard index coefficiency clustering of amino acid sequences to form VOG. A more detailed illustration is in **Additional file 1: Figure S5** and description is in Material and Method.

In order to facilitate the use of VIRGO to characterize vaginal microbial communities, each non-redundant gene was taxonomically and functionally annotated. Non-redundant genes were assigned to taxonomic groups using a custom pipeline as depicted in **Additional file 1: Figure S5.** First, metagenomic contigs were assigned taxonomy if 95% of the composite reads were annotated to the same species. Second, genes encoded on an metagenomic contig with assigned taxonomy were given the taxonomy of that contig (details in Methods). A total of 458,526 non-redundant genes comprising 48.4% of VIRGO were able to be taxonomically curated. Overall, 269 unique bacterial species were annotated in VIRGO (**Additional file 2: Table S4**), representing a majority of the described vaginal species (**Additional file 1: Figure S2**). This includes BVAB1, an as of yet unculturable vaginal species, for which several metagenome-assembled genomes (MAGs) were recently made available (accession # will be provided upon acceptance of the manuscript). BVAB1 was only been previously detectable using a partial 16S rRNA gene reference sequence [34]. It was found abundantly present in most of the metagenomes with a prevalence of 15.6% and mean abundance of (18.9% +/− 0.01) as shown in **Figure 3D**. When stratified by CST, CST IV metagenomes have the smallest proportion (<30%) of their gene content taxonomically annotated (**Additional file 1: Figure S6**) compared to ∼45-50% in *Lactobacillus*-dominated CSTs. The most abundant species based on gene content are shown in **Fig. 3a** and **Additional file 1: Figure S7.** Besides bacteria, we also curated potential fungal and phage genes (details in Methods) that were generally present in low abundance if detected at 0.17±0.04% and 0.03±0.001%, respectively. An additional 10,908 fungal and 15,965 phage genes were included (**Additional file 2: Table S5**, https://github.com/Ravel-Laboratory/VIRGO).

**Figure 3.**
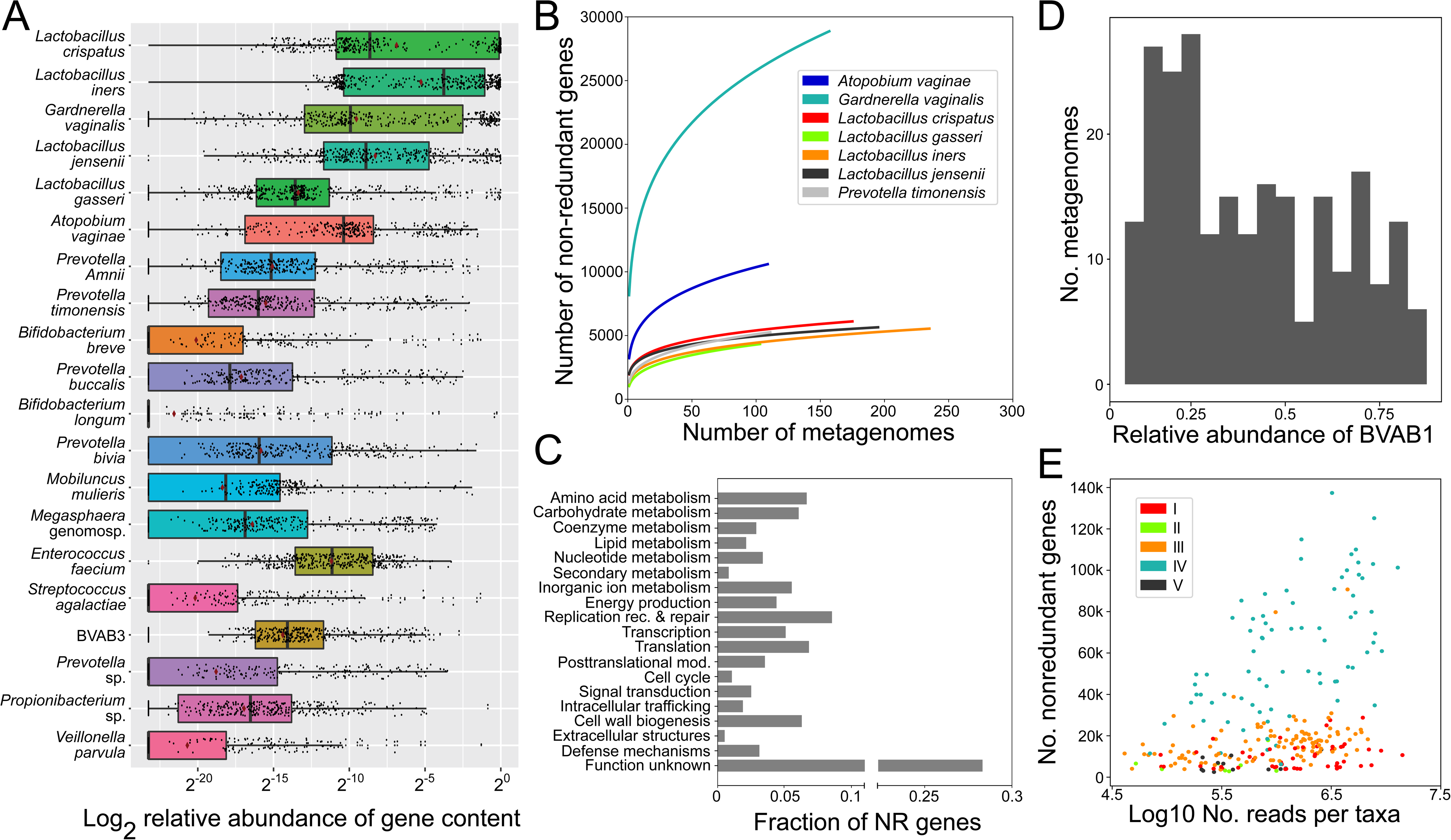
Taxonomic and functional composition of vaginal microbiome in VIRGO. (**A**) Top 20 species with the most abundant gene content in VIRGO. The logarithm of the ratio of the gene content of a species over the entire community to the base 2. Plotted are interquartile ranges (IQRs, boxes), medians (line in box), and mean (red diamond). (**B**) Species-specific metagenome accumulation curves for the number of non-redundant genes. (**C**) Functional distribution of non-redundant genes in VIRGO. Functional categories were defined using EggNOG (v4.5) [77]. (**D**) Prevalence of BVAB1 in metagenomes using a minimum number of genes threshold of 50% of the estimated BVAB1 genome size. A gene was present if ≥ 3 reads mapped to it. (**E**) Relationship between the depth of sequencing and the number of bacterial non-redundant genes identified using VIRGO. Each point is a separate metagenome and is color-coded according to community state type.

By including many metagenomes and bacterial isolate genome sequences, we sought to capture each vaginal species’ pangenome in VIRGO. To determine the extent to which we were successful, we generated metagenome accumulation curves for the number of non-redundant genes belonging to several key vaginal species (**Fig. 3b**) These curves track the number of new non-redundant genes added when increasing numbers of metagenomes containing a given species are included in constructing the database. The accumulation curves for six of the seven species tested (*L. crispatus, L. iners, L. gasseri, L. jensenii, P. timonensis, A. vaginae*) have reached saturation (**Fig. 3b**). This indicates that VIRGO includes the majority of these species pangenomes. The number of non-redundant genes included for five out of these six species are similar (∼5,000 genes), while the sixth, *A. vaginae*, had twice as many. This pales in comparison to the number of non-redundant genes included in VIRGO for *G. vaginalis*, which surpasses 25,000 genes. *G. vaginalis* is the only species analyzed for which saturation as estimated by metagenome accumulation curves, was not reached.

The non-redundant genes were decorated with a rich set of functional annotations. We performed intensive functional annotation using both the JCVI standard operating procedure [HMP] for annotating prokaryotic metagenomic shotgun sequencing data [35] as well as 17 additional functional protein databases including KEGG, COG, eggNOG, gene product, CDD, and GO, among others. A complete list of the functional annotation sources employed to characterize the VIRGO non-redundant genes is illustrated in **Fig. 2**, and an overview of the eggNOG functions encoded in VIRGO is shown in **Fig. 3c**. Overall 785,268 genes (82.8% of all non-redundant genes) were assigned a functional annotation from at least one source. This gene-rich annotation of the non-redundant gene catalog enables a comprehensive functional characterization of vaginal metagenomes and metatranscriptomes.

### VOG: orthologous protein families in vaginal microbiome

The non-redundant genes were translated into amino acid sequences and clustered into vaginal orthologous groups (VOGs). The resulting database of VOGs can be used to interrogate the protein families found in the vaginal microbiome. A modified Jaccard index was used as a measure of similarity between amino acid sequences [36, 37]. Briefly, the similarity between each pair of proteins was calculated as the intersection divided by the union of the list of proteins connected to the pair of proteins, (**Fig. 2** and **Additional file 1: Figure S5** algorithm accessible at https://github.com/Ravel-Laboratory/VIRGO). The resulting connected graph of proteins is referred as Jaccard clusters (JACs), and reciprocal best hits of JACs is referred as Jaccard orthologous clusters (JOCs) (details in Methods). The JOCs orthologous protein families can be highly conserved (alignment score >950) or partially aligned with both conserved and variable regions (alignment score ∼300) (**Additional file 1: Figure S8**). This highlights the flexibility of the network-based aggregation algorithm used to recruit both highly similar and distantly related proteins without imposing a single similarity threshold. A total of 617,127 JACs and 552,679 JOCs were generated, of which 177,684 contained at least two genes while the remaining 374,995 are singletons, indicating 38.5% of all VOG proteins are unique. The sequences, alignment, and phylogenetic trees for each of the JOCs will be available at https://github.com/Ravel-Laboratory/VIRGO.

Complementary to the VIRGO non-redundant gene sequences, VOG provides an amino acid sequence reference that can be used to improve functional annotation, comparative genomics and evolution of vaginal orthologous protein families. For example, we used VOG and retrieved 32 proteins of the orthologous family encoding vaginolysin, a *G. vaginalis* cholesterol-dependent cytolysin that is key to its pathogenicity as it forms pore in epithelial cells [38, 39] (**Additional file 2: Table S6, Additional file 1: Figure S9**). Using the retrieved alignment, we identified 3 amino acid variants in a 11-amino acid sequences of domain 4 of vaginolysin, one of the three variants, an alanine-to-valine substitution that is divergent across *G. vaginalis* and had not been reported previously. This example illustrates how VOG can be mined to understand biological relevance and to generate hypotheses. In this case it points to potential differences in pore formation activity and possibly cytotoxicity, which could be further investigated. As another example to use VOG for a large-scale data mining of protein family of interest, we searched VOG using the key phrase “cell surface-associated proteins” and “*L. iners*” and retrieved two protein families, one of which was recognized to have an LPXTG motif while the other harbored the motif YSIRK (**Additional file 2: Table S7**). Interestingly, a previous study on staphyloccocal proteins suggested that the motifs LPXTG and YSIRK were involved in different biological processes related to surface protein anchoring to cell wall envelope [40], and both are implicated in virulence by promoting bacterial attachment to alpha- and beta-chains of human fibrinogen and formation of bacterial clumps [41]. These two retrieved protein families are specific to *L. iners* and provide relevant evidence for future experimental validation to understand adherence and related biological processes. These two examples demonstrate how the VOG database can be used to explore more mechanistic understandings of vaginal bacterial communities.

### Gene richness is characteristic of vaginal microbiomes

Gene richness, calculated as number of non-redundant genes, has been adapted as the proxy of genetic diversity based on community gene content, and more recently, as community-level biomarker in gut quantitative metagenomics studies [42, 43]. We applied this paradigm to vaginal metagenomes included in VIRGO and defined high gene count (HGC) vaginal communities as those that contained >10,000 non-redundant genes and low gene count (LGC) vaginal communities as those that contained ≤ 10,000 non-redundant genes. The number of non-redundant genes identified in a metagenome was not found to correlate with the depth sequencing (**Figure 3E**. **Additional file 2: Table S8**). As expected, HGC communities had a significantly higher number of non-redundant genes (29,898±1,025) than LGC communities (4,920±151.6), however these types of communities also showed differences in their functional makeup. The LGC communities were found to be enriched for genes related to carbohydrate transport and metabolism, as well as those involved in transcription, while HGC communities were found to be enriched in genes related to intracellular trafficking, secretion, and vesicular transport, including coenzyme transport and metabolism (**Additional file 1: Figure S10**). We also found that *Lactobacillus*-dominated communities were typically categorized as LGC (82.9%) and *Lactobacillus*-deficient communities as HGC (88.3%) (**Fig. 4a**). However, this was not always the case, most notably, *L. iners*-dominated communities were classified as HGC 21.7% of the time, the highest percentage among all *Lactobacillus*-dominated communities. In fact, *L. iners-*dominated communities (7,803±6,973) generally had a greater gene richness than *L. crispatus*-dominated (5,409±3,392), *L. gasseri*-dominated (3,909±2,761), and *L. jensenii*-dominated (3,990±3,230) communities. Further, *L. iners* in HGC communities and *L. iners* in LGC communities show distinct functional makeup (**Additional file 1: Figure S11**). Similarly, not all *Lactobacillus*-deficient communities were classified as HGC—11.7% of these communities were identified as LGC. This includes communities with a high abundance of *G. vaginalis,* whose gene richness varied between 7,689±1,700 in LGC and 16,887±566 in HGC communities.

**Figure 4.**
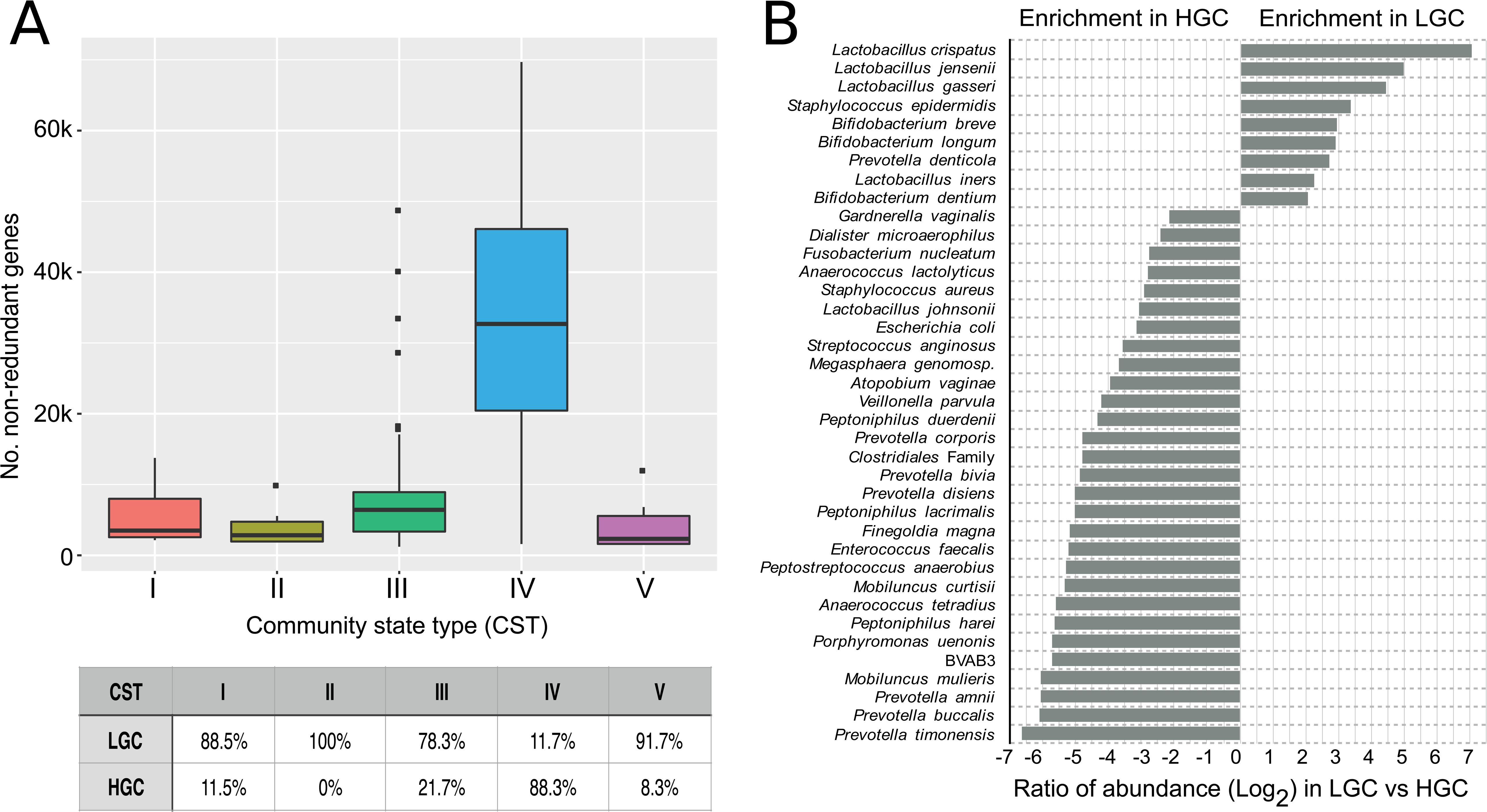
(**A**) Boxplot of the number non-redundant genes in samples of different Community State Types (CSTs). CSTs were defined as previously according to the composition and structure of the microbial community [29]. Table below boxplot contains percentage of samples in each of the CSTs stratified by high gene count (HGC) or low gene count (LGC). (**B**) Plot of the log_2_ transformed ratio of the gene of a species being in one gene count category over the other across the 264 vaginal metagenomes, only the species with more than 4 times more abundant in a category (either HGC or LGC) are shown. Plotted are interquartile ranges (IQRs, boxes), medians (line in box), and mean (red diamond).

In addition to being a characteristic of individual communities, gene richness can also be used to characterize individual genes based on their observed preference for either HGC or LGC communities. Using data from the 264 vaginal metagenomes, we classified each non-redundant gene as either an HGC or LGC gene if ≥95% its occurrences were in HGC or LGC communities, respectively. Genes that did not meet this criterion were annotated as having no preference. These gene richness annotations were included for each non-redundant gene in VIRGO. For example, 84.1%, 53.3%, 60.5% of top prevalent tryptophan biosynthesis genes in VIRGO, tryptophanase (TNAA), tryptophan synthase beta chain (TRPB), and tryptophanyl-tRNA synthetase (TRPS), are HGC genes, while 0%, 0%, and 7.0% are LGC genes (**Additional file1: Table S9**). Given the top most affiliated taxonomic groups for these tryptophan biosynthesis genes were identified as *G. vaginalis, A. vaginae, M. mulieris* (**Additional file 1: Figure S12**) our result indicates tryptophan biosynthesis genes are most prevalent in BV-associated bacteria of high gene richness vaginal communities, agreeing with recent studies [44, 45].

Using these gene annotations, we were further able to evaluate whether a vaginal bacterial species’ genes were overrepresented as being HGC or LGC (**Fig. 4b**). *Lactobacillus* spp., particularly *L. crispatus, L. jensenii, L. gasseri, L. vaginalis,* were observed to be highly overrepresented in LGC communities. On the other hand, genes belonging to many other BV-associated species, specifically *P. timonensis, P. buccalis, P. amnii, M. mulieris*, BVAB3, *Porphyromonas uenonis, P. harei, Anaerococcus tetradius, M. curtisii*, were overrepresented in HGC. These results demonstrate gene richness category information, characteristics of vaginal metagenomic communities as well as individual genes in the community, provides additional dimension to facilitate our understanding of the genetic basis of the biological processes that drive vaginal microbiomes.

### Integration of metagenome and metatranscriptome data using VIRGO as a reference framework

By serving as a reference, VIRGO enables the characterization and integrative analyses of the abundance of genes and their expression in the vaginal microenvironment. To demonstrate its use, we analyzed a woman’s vaginal metagenomes and associated metatranscriptomes at four time points over an episode of symptomatic bacterial vaginosis (BV): prior to (T1), during (T2 & T3), and after (T4) (**Fig. 5a**). Not surprisingly, the expressed functions represented in the metatranscriptomes were often different from the encoded functional makeup of the corresponding metagenomes (**Fig. 5b**). For example, T4 genes related to translation were underrepresented in the metatranscriptome as compared to the metagenome, while genes of unknown function were overrepresented. VIRGO enables rapid binning of genes by species, which revealed dramatic differences in gene abundance and their transcriptional activity in vaginal species (**Fig. 5c**). Prior to the BV episode (T1), a small proportion of *L. iners* genes were present (1.5%) but these genes exhibited high expression levels, accounting for over 20% of the metatranscriptome. At the same time point, *L. crispatus* genes made up the majority of the gene present (96.3%) but exhibited low expression levels (34.2%). In contrast, at the end of the BV episode, *L. crispatus* gene made up a small proportion of the metagenome (T3) but were highly transcriptionally active. This increased activity corresponded with *L. crispatus* regaining dominance at T4, following the resolution of the BV episode. Similarly, despite its low abundance, *P. harei* was highly transcriptionally active during the BV episode (T3), expressing transcript associated with amino acid transport and metabolism, indicating a potential role for this bacterial species in the etiology or symptomology associated with BV. Interestingly, the functional makeup of *G. vaginalis* is similar at T2 and T3, but its metatranscriptome is enriched for functions involved in energy production and conversion at T2, and enriched for functions related to translation, energy production, and carbohydrate metabolism at T3. These examples highlight how VIRGO can be used to integrate metagenome and metatranscriptomic datasets to gain better functional insights into the vaginal microbiome.

**Figure 5.**
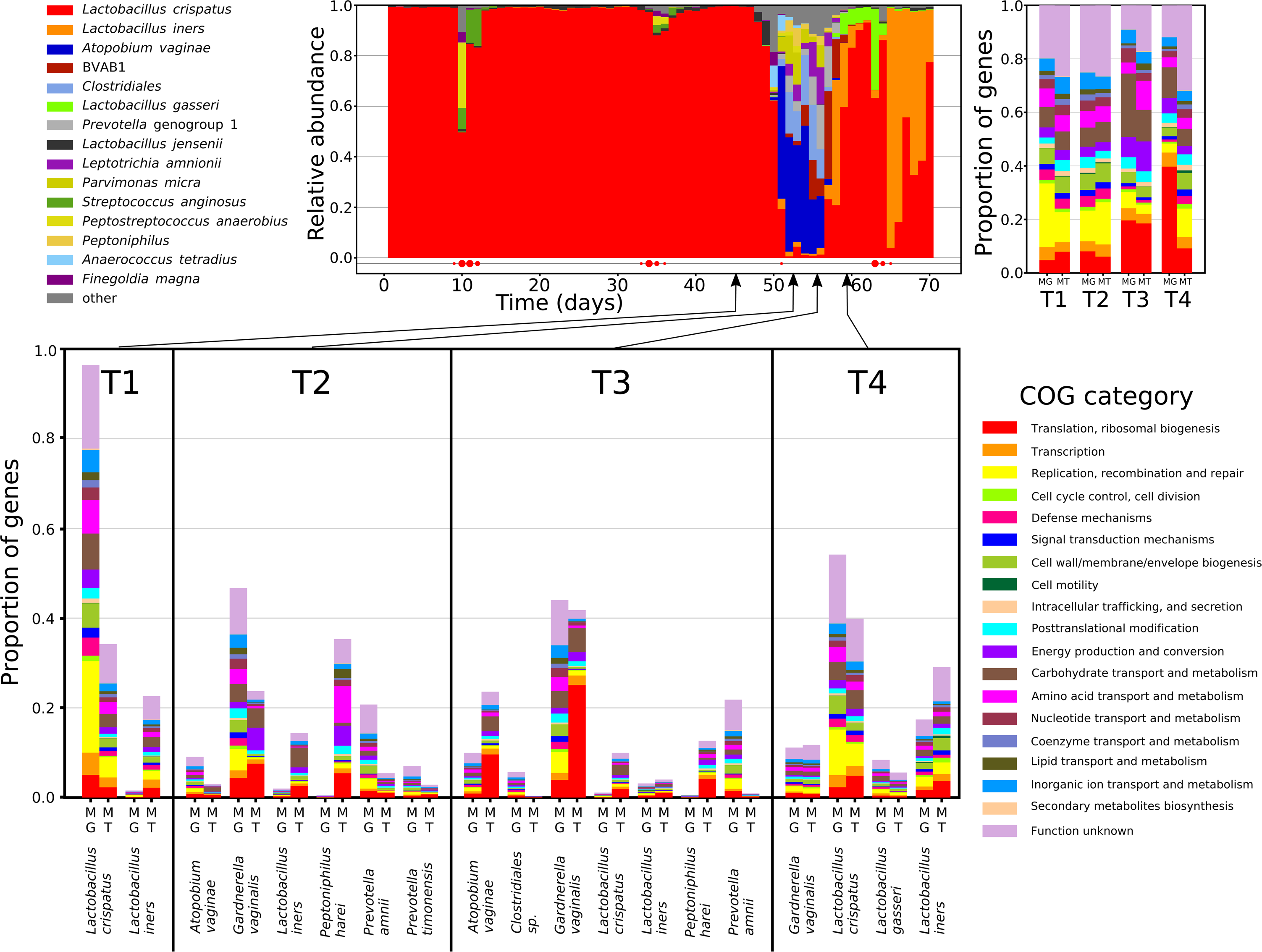
Demonstration using VIRGO and VOG to study vaginal microbiome. (**A**) 4 sampling points were selected based on a longitudinally profiled subject prior to (T1), during (T2 and T3), and after (T4) an episode of bacterial vaginosis using 16S rRNA profiling. (**B**) Functional profiling of the metagenome (MG) and metatranscriptome (MT) of each of the 4 sampling points. Functional categories were annotated using EggNOG (v4.5) [77]. (**C**) Functional profiles stratified by species using the taxonomic profiling provided by VIRGO. (**D**) Demonstrative use of VOG to characterize the *G. vaginalis* cholesterol-dependent cytolysin (CDC) protein family. It shows the phylogeny of CDC-containing protein and alignment of domain 4 of the CDCs that is generally well conserved but contains a single divergent site, highlighted in yellow [38].

### VIRGO revealed high within-community intraspecies diversity

VIRGO can be used to characterize the genome content of individual bacterial species that are present in the vaginal microbiome. We applied VIRGO to a dataset of 1,507 *in-house* and publicly available vaginal metagenomes, to characterize the gene content of four *Lactobacillus* species (*L. crispatus, L. iners, L. jensenii,* and *L. gasseri)* and three additional species commonly found in the vagina (*G. vaginalis, A. vaginae* and *P. timonensis*). We recovered most of each species gene content (>80% of the average gene count in a genome) even when that species was present at low abundance (<1%) in a community. For instance, even though *P. timonensis* [46] was generally present in low abundance in these metagenomes (4.8% ± 0.3% mean ± S.E., range [0.1-33.8%]), we recovered the majority of its genome (2,469±401 CDS, **Additional file 1: Figure S13; Additional file 2: Table S10**). We observed similarly high sensitivity in the analysis of the other six selected vaginal species (**Fig. 6a, Additional file 2: Table S10**). These results demonstrate VIRGO’s capability for characterizing the gene content of low abundance taxa from metagenomic data.

**Figure 6.**
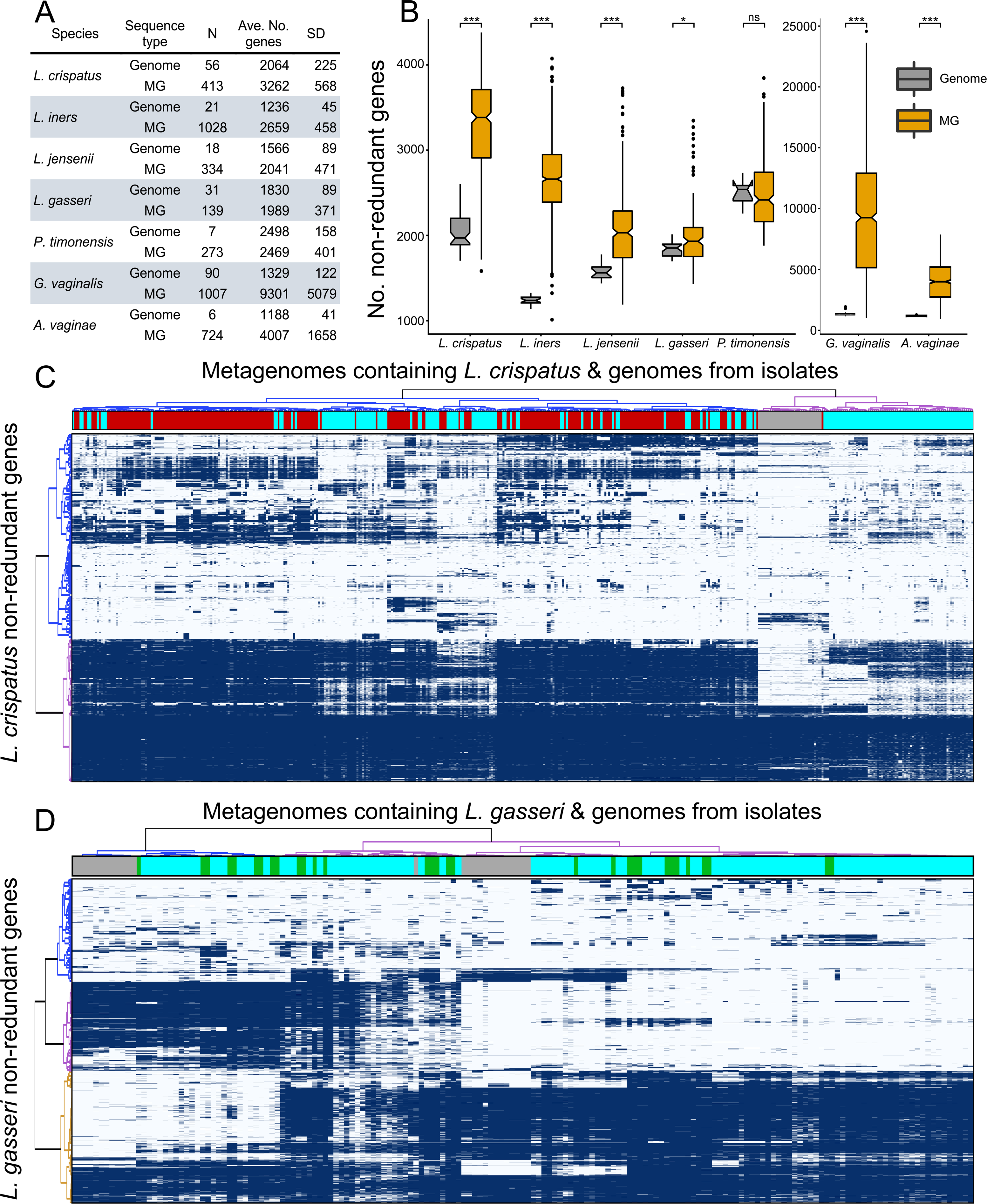
Intraspecies diversity revealed using VIRGO of seven vaginal species including *L. crispatus, L. iners, L. jensenii, L. gasseri*, and *G. vaginalis, A. vaginae* and *P. timonensis*. (**A**) Summary of the number (N) of isolate genomes and metagenome (MG) samples with more than 80% of their average genome’s number of coding genes for a species, based on a dataset of 1,507 *in-house* vaginal metagenomes characterized using VIRGO. (**B**) Boxplot of number non-redundant genes in isolate genomes versus vaginal metagenomes. (**C**) Heatmap of presence/absence of *L. crispatus* non-redundant gene profiles for 56 available isolate genomes and 413 VIRGO-characterized metagenomes that contained either high (red) or low (blue) relative abundance of the species. Hierarchical clustering of the profiles was performed using ward linkage based on their Jaccard similarity coefficient. * number of isolate genomes and metagenome samples. ^†^ MG: Metagenomes *p<0.05,***p<0.001 after correction for multiple comparisons.

Using these species-specific gene repertoires, we characterize the amount of intraspecies diversity present within an individual woman’s vaginal microbiome. Because VIRGO basically comprises the “pangenomes” of each vaginal bacterial species, it can be used to evaluate the amount of intraspecies diversity present in these communities. For this analysis, we counted the number of genes that were assigned to each of the seven species in each of the 1,507 metagenomic datasets and compared this number to that found in each species’ reference genomes. The number of genes for a species in a community often exceeded that found in a single isolate genome (**Fig. 6a, 6b**), suggesting that multiple strains of a species co-occur in vaginal bacterial communities. The total number of *L. crispatus* genes identified in each of the metagenomes where it was detected contained on average 1.6 times more genes (3,262±586) than that found encoded on *L. crispatus* genomes (2,064±225, P<0.001). Similar results were observed for *G. vaginalis, A. vaginae, L. iners, L. jensenii*, and *L. gasseri*, which are represented by 7.0, 3.4, 2.2, 1.3, and 1.1 times more genes in metagenomes than that found in genomes, respectively. *G. vaginalis* and *A. vaginae* exhibited the highest degree of intraspecies diversity, while *L. crispatus* has the highest within-metagenome intraspecies diversity among all major vaginal *Lactobacillus* spp. (**Additional file 1: Figure S13; Figure 6c**). These results suggest that a woman’s vaginal bacterial populations are routinely comprised of more than one strain of most species. VIRGO affords investigating this unprecedented intraspecies diversity in vaginal communities.

We next applied well-established practices from pangenomics [10, 12] in order to identify core and accessory non-redundant genes among our sample-specific species gene repertoires. Based on the clustering patterns of gene prevalence profiles, we were able to define groups of consistently present (core) and variably present (accessory) non-redundant genes. The majority of the observed genes for each of the species were categorized as accessory, with variable representation across the metagenomic datasets. Using *L. crispatus* as an example, we observed more than twice as many non-redundant genes with variable representation across the metagenomes than those present in every sample (**Fig. 6c**). Interestingly, it is clear from this analysis that the gene content identified with VIRGO in genome sequences of *L. crispatus* under-represent the intraspecies genetic diversity present in the metagenomes. Similar results were observed for the other six species analyzed, although the magnitude of the difference between the metagenome and isolate gene repertoires varied depending on the species. Overall, VIRGO revealed that metagenomic data carry a more extensive gene content than is found in all combined isolate genome sequences.

### Metagenomic subspecies in vaginal ecosystem

Hierarchical clustering of the metagenome species-specific gene content profiles revealed distinct groupings which we term “metagenomic subspecies” (MG-subspecies). These metagenomic subspecies represent types of bacterial populations that share a similar gene pool as assessed by shotgun metagenomic sequence data. For example, this analysis revealed at least three distinct metagenomic subspecies for *L. gasseri* (**Fig. 6d**). *L. gasseri* MG-subspecies I and III have large sets of non-redundant genes that are present in one but not the others, while *L. gasseri* MG-subspecies II carries a blend of the genes from both MG-subspecies I and III. The analysis of *G. vaginalis* revealed more than four types of profile groupings, though concordant with the previously described multiple types of isolate genomes [47], we find that this genome-based paradigm largely under-represents the diversity of *G. vaginalis* gene content identified in metagenomes (**Additional file 1: Figure S13e**). We applied this analysis to seven vaginal species (**Additional file 1: Figure S13**) and found that vaginal microbial communities are often composed of complex mixtures of multiple strains of the same species, and that these mixtures can be clustered into distinct MG-subspecies. Further interrogation of these vaginal MG-subspecies and their gene content is likely to reveal novel features of vaginal communities and their sub-populations that will contribute to our understanding of the vaginal ecosystem of niche-optimized strains.

## Discussion

Microbiome studies have become increasingly sophisticated with the rapid advancement of sequencing throughput and the associated decrease in sequencing cost. However, identifying features that drive correlations between the microbiome and health using multi-omics sequence data remains challenging. This is due, in part, to difficulties in analyzing and integrating the complex, feature rich, metagenomic and metatranscriptomic data now common to microbiome studies. A scalable tool that provides a comprehensive characterization of such multi-omics data is therefore highly desired. VIRGO is a large vaginal microbiome database designed to fulfill such research needs for investigations of the vaginal microbiome and its relation to women’s health. In summary, VIRGO has (i) a comprehensive breadth that includes previously observed community types, vaginal species, and even fungi and viruses; (ii) a gene-centric design that enables the integration of functional and taxonomic characterization of metagenomic and metatranscriptomic data originating from the same sample; (iii) a high scalability and low memory requirement; (iv) a high sensitivity that affords characterization of the gene content of low-abundance bacteria; (v) an easy to use framework from which to evaluate gene richness and within-species diversity.

VIRGO contains a multitude of non-redundant genes that we identified in vaginal metagenomes and urogenital bacterial isolates. These non-redundant genes were also clustered into orthologous groups (VOGs) using a memory-efficient network-based algorithm that handles nodes connectivity in high dimensionality space [48, 49]. This approach to identifying orthologous protein sequences allows for great flexibility because it does not rely on a single sequence similarity cutoff value [50, 51]. These families of vaginal orthologs will assist the development of a mechanistic understanding of these proteins and how they relate to health. For example, van der Veer and co-workers recently identified and characterized the *L. crispatus* pullulanase (*pulA*) gene which they show encodes an enzyme with amylase activity that likely allows this species to degrade host glycogen in the vaginal environment [52]. Using VIRGO and VOG, we were able to identify pullulanase domain containing proteins in 37 other vaginal taxa including: *G. vaginalis, L. iners* and *P. timonensis* (**Additional file 2: Table S12**), providing insight into the breadth of vaginal bacteria that may be capable of degrading host glycogen. In this way, VIRGO and VOG can facilitate knowledge retrieval, hypothesis generation and future experimental validation to advance understanding of vaginal ecosystem.

Using VIRGO, we observed that vaginal metagenomes varied in gene richness, with some communities having more non-redundant genes than others. Gene richness has been found to be indicative of the pathophysiological state of the gut microbiome in studies of obesity [43], dietary intervention [42], type II diabetes [53], and inflammation and metabolic disease [54]. We adapted the concept of gene richness as a characterization of community gene content and defined an analogous definition for the vaginal microbiome. An outstanding difference in gene richness was observed between *Lactobacillus*-dominated and *Lactobacillus*-deficient communities. Approximately 85% of communities with a high relative abundance of *Lactobacillus* sp., had a low gene richness across the community, whereas *Lactobacillus*-deficient communities were more likely to have a high gene richness. However, around 22% of *Lactobacillus*-dominated communities did have high gene richness and 12% of *Lactobacillus*-deficient communities had low gene richness. It may be that gene richness category, when combined with community state types, provides a useful and, ecologically relevant, categorization of vaginal community states. For example, it is envisioned that a subject that has a *Lactobacillus*-dominated community with high gene richness is at a higher risk of switching to a dysbiotic state than one whose community is dominated by *Lactobacillus* but with low gene richness. In such case, VIRGO provides the analytical suite needed to test this and other hypotheses relating gene richness to the ecology of the vaginal microbiome.

In our demonstrative analysis of more than 1,500 metagenomes, we identified and characterized a wealth of intraspecies diversity that was present within individual vaginal microbial communities. Populations of bacterial species in vaginal communities comprises of multiple strains. Previous studies of the vaginal microbiome have largely treated these species as singular genotypes [55, 56], although some more recent studies have examined intraspecies diversity in these communities [57, 58]. Intraspecies diversity is important because it is likely to influence many properties of the communities including their temporal stability and resilience, as well as how they relate to host health. Unfortunately, intraspecies diversity is difficult to detect using typical assembly-based metagenomic analysis strategies, which are notoriously ill suited for resolving strains of the same species [59, 60]. VIRGO can be a more suitable tool for characterizing intraspecies diversity because it was built to contain the non-redundant pangenomes of most species common to the vagina. Strict mapping of sequence reads against the VIRGO database provides an accurate and sensitive way of identifying the aggregated non-redundant genes that belongs to each species in a metagenome. We expect VIRGO to facilitate future investigations of intraspecies diversity in vaginal microbial communities. We further showed that, for the seven species we examined, the intraspecies diversity had structure. Vaginal metagenomes from different subjects contained related sets of species-specific non-redundant genes. We postulate that these clusters of samples with shared gene content represent similar collectives of strains which we have termed “metagenomic subspecies”. It is expected that, given their shared gene content, these metagenomic subspecies might also share phenotypic characteristics. However, additional studies are needed to characterize differences between metagenomic subspecies and to detail their possible effect on host health. Reconstructing a particular species’ metagenomic subspecies might be possible by identifying and combining isolates that adequately cover the genetic repertoire of the metagenomic subspecies. One complication to this approach is that, in many cases, the observed metagenomic subspecies contain non-redundant genes which have not been observed in isolate genome sequences for that species. This could reflect a limitation in the number of isolate genomes available for a species or even systematic bias in the growth and recovery of species from the vagina [61]. Targeted isolation of strains from communities containing the metagenomic subspecies of interest are needed in order to fill in these gaps in the future.

The value of VIRGO resides in its functions as both a central repository and a highly scalable tool for fast, accurate characterization of vaginal microbiomes. VIRGO is particularly useful for users with limited computational skills, a large volume of sequencing data, and/or limited computing infrastructure. In particular, the metagenome-metatranscriptome data integration enabled by the gene-centric design in VIRGO provides a powerful approach to determine the expression patterns of microbial functions, and in doing so, to characterize contextualized complex mechanisms of host-microbiota interactions in vaginal communities. This feature makes possible the meta-analyses of vaginal microbiome features and the quantitative integration of findings from multiple studies, which helps with the common issue of confounding gene copy number that has been a major challenge in analyzing metatranscriptomic dataset [62, 63]. We also anticipate that VIRGO will be used to process metaproteomic datasets when that practice becomes common and easily accessible. Each of the protein sequence of each gene could be used to map peptides obtained from metaproteomic pipelines and access VIRGO rich annotation. On the other hand, we acknowledge the limitations of the referenced based approach of VIRGO. This version is focused on the gene-level de-redundancy and characterization of vaginal microbiome. However, in the future we plan to expand VIRGO to include the capability to identify nucleotide variants within a gene. We believe this will further facilitate our understanding of within-species diversity and evolutionary change in the vaginal ecosystem. The database is primarily focused on bacteria with limited inclusion of viral and fungal gene sequences. Future in-depth profiling of these non-bacterial microbes will allow VIRGO to provide a more complete picture of vaginal microbial communities.

## Conclusion

Efforts are underway to translate our growing understanding of human-associated microbial communities into clinical biomarkers and treatments. A deeper understanding of the complex mechanisms of host-microbiota interactions requires the integration of multi-omics data. VIRGO presents a central reference database and analytical framework to enable the efficient and accurate characterization of the microbial gene content of the human vaginal microbiome. Powered by a rich suite of functional and taxonomic annotations, VIRGO allows for the integrated analysis of metagenomic and metatranscriptomic data. VIRGO further provides a gene-centric approach to describe vaginal microbial community structure including fine scale variation at the intraspecies level. This unprecedented view of intraspecies diversity within a vaginal community is far beyond the scope offered by current genome references. VIRGO is a centralized, and freely available resource for vaginal microbiome studies. It will facilitate the analysis of multi-omics data now common to microbiome studies, and provide comprehensive insight into community membership, function, and ecological perspective of the vaginal microbiome.

## Methods

### Datasets

Metagenomes used in this study include 211 newly *in-house* sequenced datasets and 53 vaginal datasets downloaded from the HMP data repository (http://www.hmpdacc-resources.org/cgi-bin/hmp_catalog/main.cgi). Genome sequences of urogenital bacterial isolates deposited in multiple databases were downloaded on November 10, 2016, including GenBank (http://www.ncbi.nlm.nih.gov/), IMG/M: Integrated Microbial Genomes & Microbiomes (https://img.jgi.doe.gov/), and HMP referencing genome database (http://www.hmpdacc-resources.org/cgi-bin/hmp_catalog/main.cgi). After removing duplicate genomes under the same strain names, genomes of 416 urogenital bacterial strains and 321 bacterial species were included in the catalog. A full list of the genomes and metagenomes used in the construction of the database can be found in **Additional file: Table S1**.

### Nucleic acid extraction, library construction, and metagenome and metatranscriptome sequencing

The included 211 *in-house* metagenomes were generated as follows: whole genomic DNA was extracted from 300 µl aliquot of vaginal ESwab re-suspended into 1ml Amies transport medium (ESwab, Copan Diagnostics Inc.) and preserved at −80°C. Briefly, Cells were then lysed using a combination of enzymatic digestion and mechanical disruption that included mutanolysin, lysostaphin and lysozyme treatment, followed by proteinase K, SDS and bead beating steps. Procedures for DNA extraction and concentration qualification were previously described [29, 64]. The shotgun metagenomic sequence libraries were constructed from the extracted DNA using Illumina Nextera XT kits and sequenced on an Illumina HiSeq 2500 platform at the Genomic Resource Center at the University of Maryland School of Medicine.

The metatranscriptomes used to demonstrate the use of VIRGO for the analysis of community-wide gene expression were obtained from RNA extracted from vaginal swabs stored in 2 ml Amies Transport Medium-RNAlater solution (50%/50%, vol/vol) archived at −80°C. A total of 500 µl of ice-cold PBS was added to 1,000 µl of that solution and spun down at 8,000x*g* for 10 min. The pellet was resuspended in 500 µl ice-cold RNase-free PBS with 10 µl β-mercaptoethanol. The suspension was transferred to Lysis Matrix B tube (MP Biomedicals) containing 100 µl 10% SDS and 500 µl acid phenol and beads beaded using a FastPrep instrument (MP Biomedicals) for 45 seconds at 5.5 m/s. The aqueous phase was mixed with 250 µl acid phenol and 250 µl 24:1 chloroform:isoamyl alcohol. The aqueous layer was again transferred to a fresh tube and mixed with 500 µl 24:1 chloroform:isoamyl alcohol. For every 300 µl resulting aqueous solution, we added 30 µl of 3 M sodium acetate, 3 µl of glycogen (5 mg/ml), and three volumes of 100% ethanol. The mixture was incubated at −20°C overnight to precipitate the nucleic acids. After centrifugation at 13,400x*g* for 30 min at 4°C, the resulting pellet was washed, dried, and dissolved in 100 µL of DEPC-treated water. Carryover DNA was removed by: 1) treating twice with Turbo DNase free (Ambion, Cat. No. AM1907) at two half-hour intervals according to the manufacturer’s protocol for rigorous DNAse treatment, 2) purifying twice using gDNA-eliminator columns (QIAGEN) before and after DNase treatment followed by RNeasy column purification (QIAGEN). We further conducted PCR using 16S rRNA primer 27F (5’-AGAGTTTGATCCTGGCTCAG-3’) and 534R (5’-CATTACCGCGGCTGCTGG-3’) to confirm DNA removal. The quality of extracted RNA was checked using an Agilent 2100 Expert Bioanalyzer Nano chip. Ribosomal RNA removal was performed according to the manufacturer’s protocol of a combined Gram-positive, Gram-negative and Human/mouse/rat Ribo-Zero rRNA Removal Kit (Epicentre Technologies). The resulting RNA was purified using Zymo RNA clean & Contentrator-5 column kit (ZYMO Research). RNA final quality was checked using an Agilent RNA 6000 Expert Bioanalyzer Pico chip. Sequencing libraries of A and B containing 6 bp indexes were prepared using the TruSeq RNA sample prep kit (Illumina) following a modification of the manufacturer’s protocol: cDNA was purified between enzymatic reactions and library size selection was performed with AMPure XT beads (Beckman Coulter Genomics). Library sequencing was performed using the Illumina HiSeq 2500 platform.

### Construction of the human vaginal non-redundant gene catalog (VIRGO)

Multiple bioinformatics pre-processing steps were applied to the raw shotgun metagenomic sequence datasets, including (1) eliminating all human sequence reads (including human rRNA LSU/SSU sequence reads) using BMTagger v3.101 [65] against a standard human genome reference (GRCh37.p5 [66]), (2) *in silico* microbial rRNA sequence reads depletion by aligning all reads using Bowtie (v1) [67] against the SILVA PARC ribosomal-subunit sequence database [19] to eliminate mis-assemblies of these repeated regions. After each of these steps, the paired reads were removed; (3) stringent quality control using Trimmomatic [68], in which the Illumina adapter was trimmed and reads with average quality greater than Q15 using a sliding window of 4 bp with no ambiguous base calling were retained. MetaPhlAn (v2) [69] was subsequently used to establish taxonomic profiles after these pre-processing steps. Samples were then clustered in community state types (CSTs) using taxa abundance tables and the Jensen-Shannon divergence metrics as previously described [29, 70]. Species accumulation curves and diversity estimates for rarefied samples were computed using R package *iNEXT* [71] and *vegan* [72]. The 264 vaginal metagenomes were then assembled using IDBA-UD (v1.0) [73] with a k value range of 20-100. Genes were called on the resulting contigs using MetageneMark (v3.25) [33] to predict CDSs with the default settings. Genes and gene fragments that were at least 99bp long, with greater than 95% identity over 90% of the shorter gene length were clustered together by a greedy pairwise comparison implemented in CD-HIT-EST (v4.6) [74], according to the clustering procedure and threshold defined previously [16, 22]. The gene with the longest length ≥99bp was used as the representative for each cluster of redundant genes.

### Taxonomic and functional annotations of VIRGO

The non-redundant genes were annotated with a rich set of taxonomic and functional information. Genes that originated from an isolate sequence genome were automatically assigned that species name. For metagenomes, taxonomy was assigned to a metagenomic contig by mapping the sequence reads making up that contig to the Integrated Microbial Genomes (IMG) reference database (v400) using bowtie (v1, parameters: “-l 25 --fullref --chunkmbs 512 --best --strata -m 20”). A secondary filter was applied so that the total number of mismatches between the read and the reference was less than 35, and that the first 25 bp of the read matched the reference. Using the results of this mapping, taxonomy was assigned to all genes encoded on the contig that met the following four criteria: 1) at least 95% of the reads mapped to the same bacterial species, 2) the remaining 5% off-target reads did not map to a single species, 3) the contig had at least 2X average coverage and >50 reads, 4) at least 25% of the contig length had reads mapped onto. These stringent criteria were used to ensure high fidelity of the taxonomic assignments and a low contribution of potentially chimeric contigs. To further diminish the risk of incorporating false taxonomic assignments, the annotations of the contigs belonging to species at low relative abundance in the sample were removed. Genome completeness was estimated as the fractional representation of the genome in the metagenome using BLASTN (minimal overlapping >60% of the shorter sequence and >80% sequence similarity). For each metagenome, only taxonomic assignments originating from species with at least 80% representation were incorporated. The genes that shows >80% sequence similarity over 60% of query gene length to the non-redundant genes were then assigned. The non-redundant genes in VIRGO were searched against fungal database that includes 5 vaginal yeast species in 40 genomes (listed in **Additional file 2: Table S5**) using BLASTN, that a gene must have at least 80% sequencing similarity with over 60% overlapping length to be curated. We also annotated potential phage genes that may be present in VIRGO by searching against phage orthologous groups or Prokaryotic virus orthologous groups (version 2016) [51, 75], using BLASTN and included the ones at >80% sequence similarity over 60% of query gene length in annotation (**Additional file 2: Table S5**). Functional annotations based on the standard procedure for each of 17 functional databases, including: cluster of orthologous groups (COG[76], eggNOG (v4.5) [77], KEGG[78]), conserved protein domain (CDD[79], Pfam[80], ProDom[81], PROSITE[82], TIGRFAM[83], InterPro[84]), domain architectures (CATH-Gene3D[85, 86], SMART[87]), intrinsic protein disorder (MobiDB[88]), high-quality manual annotation (HAMAP[89]), protein superfamily (PIRSF[90]), a compendium of protein fingerprints (PRINTS[91]), and gene product attributes (Gene Ontology [92], JCVI SOP [35]).

### Construction of vaginal orthologous groups (VOGs) for protein families

The non-redundant genes were also clustered based on orthology to generate a set of Vaginal Orthologous Groups (VOGs). To do this we used a modified version of a Jaccard clustering method previously implemented [36, 37]. We performed an all-versus-all BLASTP search among the translated coding sequences (CDS) of the non-redundant genes included in VIRGO [93, 94]. The all-against-all BLASTP matches was used to compute Jaccard similarity coefficient for each pair of translated CDSs, without constraints based on which sample or microorganism from which it originated. Only BLASTP matches with 80% sequence identity and 70% overlap, and an E-value less than 1E-10 were used in the calculation of the Jaccard similarity coefficient. The filtered BLASTP results were then used to define connections between pairs of translated CDSs resulting in a network graph with the translated CDSs as nodes and their connections as edges. The Jaccard similarity coefficient was then calculated as the number of nodes that had direct connections to the two translated CDSs divided by the total number of nodes that had direct connections to either of the two translated CDSs in the network (intersection and union) [37]. Jaccard clusters (JACs) were defined as a set of translated CDSs whose Jaccard similarity coefficient was at least 0.55. If two translated CDSs from different JACs were reciprocal best matches according to the BLASTP searches, the two JACs were merged. Finally, the alignment program T-Coffee [95] was used to assess the alignment quality within the JACs and to calculate the alignment score.

### Bioinformatics analysis

The comprehensiveness of VIRGO was tested using vaginal metagenomic data from vaginal metagenomes of North American women not including in the construction of VIRGO and sequenced in this study, as well as women from African [30], and China [31]. The sequences reads were first mapped to the VIRGO contigs using bowtie (v2; parameters: --threads 4 --sensitive-local -D 10 -R 2 -N 0 -L 22 -i S,1,1.75 -k 1 --ignore-quals --no-unal) [96], according to the criteria used previously in the construction of a gut gene catalog [16]). Any unmapped reads were compared to the GenBank nt database [97] using BLASTN and an E-value of 1E-10 as cutoff. To annotate BVAB1 genes in VIRGO, we used BLASTN and an E-value of 1E-10 as cutoff, the matched genes with percent identity >95% over >90% of gene length were annotated as BVAB1 genes. To retrieve pullulanase (*pulA*) genes in VIRGO, we used conserved protein domain CDD [79] annotation and keyword “pullulanase”. To further demonstrate the comprehensiveness of VIRGO and that VIRGO captures the pangenome of selected species, species specific metagenome accumulation curves for the number of non-redundant genes were constructed for seven vaginal species by rarefaction with 100 bootstraps: *L. crispatus, L. iners, L. jensenii, L. gasseri*, and *G. vaginalis, A. vaginae* and *P. timonensis*.

For gene count category and analysis, the included 264 vaginal metagenomes were classified as either having a high gene count (>10,000 non-redundant genes) or low gene count (<10,000 non-redundant genes). The VIRGO non-redundant genes were then annotated as either being a high gene count gene or low gene count gene if the gene was preferentially identified (at least 95%) in high or low gene count metagenomes. The log ratio of genes of a species being in either high or low gene count metagenomes across the 264 vaginal metagenomes was calculated for all species with at least 0.1% abundance and at least 100 genes in either HGC or LGC groups. The species with more than 4 times more abundant (in logarithm 2 scale) in a category (either HGC or LGC) were considered showing preference in one of the categories.

### Using VIRGO to characterize within community intraspecies diversity

Intraspecies diversity analyses were conducted by mapping isolate genome sequences as well as vaginal metagenomes to VIRGO. We chose to focus our analysis on the previously mentioned seven vaginal species. Accession numbers for genomes of the four *Lactobacillus* species (*L. crispatus, L. iners, L. jensenii,* and *L. gasseri)* and three additional species (*G. vaginalis, A. vaginae* and *P. timonensis*) can be found in **Additional file 2: Table S11**. A total of 1,507 vaginal metagenomes including 1,403 *in-house* from de-identified vaginal swab and lavage specimens and 76 publicly available, were mapped against VIRGO. Their accession numbers can be found in **Additional file 2: Table S12**. For each of the seven species, a presence/absence matrix for the species’ non-redundant genes was constructed that included the data from species’ isolate genomes and all metagenomes that contained at least 80% of the average number of genes encoded on a genome of that species. Comparisons of the number of non-redundant genes present in the species isolate genomes versus the metagenomes in which they appeared where conducted using student t-test. Hierarchical clustering was performed on the boolean matrix of the species’ non-redundant genes using Jaccard clustering implemented in the *vegan* package in R [98]. A tutorial describing how to use VIRGO and VOG is available online at https://github.com/Ravel-Laboratory/VIRGO.

## List of abbreviations

VIRGO: human vaginal non-redundant gene catalog
VOG: vaginal orthologous groups
IMG/M: Integrated Microbial Genomes & Microbiomes
rRNA: ribosomal ribonucleic acid
CST: vaginal community state type
MAG: metagenome-assembled genome
JAC: Jaccard cluster
JOC: Jaccard orthologous cluster
HGC: high gene count
LGC: low gene count
MG-subspecies: metagenomic subspecies
Av: A. vaginae
Gv: G. vaginalis
Pt: P. timonensis
Lc: L. crispatus
Li: *L. iners*
Lj: *L. jensenii*
Lg: *L. gasseri*

## Declarations

### Electronic supplementary material

**Additional file 1: Figure S1.** Boxplot of the proportion of sequencing reads after removing human contaminates from the samples between different Community State Types (CSTs). CSTs were defined as previously according to the composition and structure of the microbial community [29]. Plotted are interquartile ranges (IQRs, boxes), medians (line in box), and mean (red diamond). Significance value was calculated using Wilcoxon rank sum test using *ggsignif* R package [99]. Star sign (*) denotes the level of significance.

**Additional file 1: Figure S2.** Heatmap of relative abundance of the 50 most abundant phylotypes in the vaginal metagenomes used in this study. Ward linkage clustering is used to clusters samples based on their Jensen-Shannon distance calculated in the *vegan* package in R [100] according to the previous naming convention [29]. The sidebars indicate CSTs and gene richness category, respectively. Gene richness categories include high gene count (HGC) and low gene count (LGC), defined using the threshold of 10,000 genes per sample.

**Additional file 1: Figure S3.** Vaginal community accumulation curves and diversity estimate. (A) Accumulative diversity estimates with respect to sample size, for rarefied and extrapolated estimates using all samples; (B) accumulative diversity estimates with respect to sample size, for rarefied and extrapolated estimates using samples of different CSTs; (C) diversity estimate with respect to sample coverage, for rarefied and extrapolated estimates using all samples; (D) diversity estimate with respect to sample coverage, for rarefied and extrapolated estimate using samples of different CSTs. Community diversity estimates were computed using R package *iNEXT* [71] and *vegan* [72]. Sampling curve was either rarefied to smaller sample sizes or extrapolated to a larger sample size for species diversity estimate.

**Additional file 1: Figure S4.** Pie chart taxonomic distribution of reads that failed to map on VIRGO for vaginal metagenomes of African women from Gosmann *et al.* [30] in **A** and of Chinese women from [31] in **B**. The unmapped reads were compared to GenBank nt database [97] using BLASTN.

**Additional file 1: Figure S5.** Pipeline for data processing and integration for the construction of the human <u>v</u>aginal integrated non-<u>r</u>edundant <u>g</u>ene catal<u>o</u>gue (VIRGO) and vaginal orthologous protein family groups (VOG). Metagenomes from 264 vaginal metagenomes and 416 genomes of urogenital isolates were processed, that including 212 *in-house* sequenced vaginal metagenomes. The procedures include pre-processing to remove human contaminates, quality assessment, metagenome assembly, gene calling, functional and taxonomic annotation, gene clustering based on nucleotide sequencing similarity to form VIRGO, and Jaccard index coefficiency clustering of amino acid sequences to form VOG.

**Additional file 1: Figure S6.** Proportion of the assembly length assigned taxonomically from the samples (**A**) among different community state types (CSTs) and (**B**) between different gene richness category. CSTs were defined as previously according to the composition and structure of the microbial community [29]. Gene richness category includes high gene count (HGC) and low gene count (LGC), defined using the threshold of 10,000 genes per sample.

**Additional file 1: Figure S7.** Top 20 species with the most abundant gene content in VIRGO. The ratio of the gene content of a species over the entire community to the base 2. Plotted are interquartile ranges (IQRs, boxes), medians (line in box), and mean (red diamond).

**Additional file 1: Figure S8.** Boxplot of the alignment scores of Jaccard orthologous clusters (JOCs) with multiple members. The alignment program T-Coffee [95] was used to access the alignment quality using alignment score.

**Additional file 1: Figure S9.** Phylogeny that is demonstrative use of VOG to characterize the *G. vaginalis* cholesterol-dependent cytolysin (CDC) protein family. It shows the phylogeny of CDC-containing protein and alignment of domain 4 of the CDCs that is generally well conserved but contains a single divergent site, highlighted in yellow [38].

**Additional file 1: Figure S10.** Association plot of functional distribution of different gene count categories in vaginal microbiome. Functional category was defined using EggNOG (v4.5) [77] functional category. A Cohen-Friendly association plot [101, 102] was produced in statistical package *vcd* in R [103] to indicate deviations to indicate deviations from independence of CSTs and functional distribution. Mosaics display was shown, where the cells are shaded in proportion to standardized residuals, where the positive value (blue) is the observed frequency is substantially greater than would be found under independence, and the negative value (red) indicates cells which occur less often than under independence.

**Additional file 1: Figure S12.** Functional category of *L. iners* in different gene richness categories. Functional category was defined using EggNOG (v4.5) [77] functional category.

**Additional file 1: Figure S13.** Gene richness category and taxonomic distribution of tryptophan production-related genes in VIRGO. (**A**) Pie chart of the percentage of tryptophan production-related genes in different gene richness categories of HGC or LGC. (**B**) The top 10 most affiliated taxonomic groups of the tryptophan production-related genes.

**Additional file 1: Figure S14.** Heatmap includes gene prevalence profiling of available genomes of vaginal isolates and VIRGO-characterized metagenomes for (**A**) *L. crispatus*, (**B**) *L. iners*, (**C**) *L. jensenii*, (**D**) *L. gasseri*, and (**E**) *G. vaginalis*, (**F**) *A. vaginae* and (**G**) *P. timonensis*. Hierarchical clustering of the profiles was performed using ward linkage based on Jaccard similarity coefficient. CSTs were defined as previously according to the composition and structure of the microbial community [29].

**Additional file 2: Table S1**. Statistics of the sequence reads, including 211 *in-house* sequenced metagenomes, 53 metagenomes from HMP DACC database, 277 genomes isolated from vagina, reproductive or urinary system deposited in GenBank and 139 urogenital bacteria genomes from HMP DACC database used to compile database. The assembly statistics includes assembled base pairs, total number of contigs, N50, mean and median length, and other statistics.

**Additional file 2: Table S2**. OTUs table for all metagenomes included in VIRGO. Taxonomic profiling was conducted in MetaPhlAn version 2 [69]. Community state types were defined as previously according to the composition and structure of the microbial community [29]. 312 bacterial species that were present in at ≥ 0.01% relative abundance are shown.

**Additional file 2: Table S3**. Statistics of the complete and subsets of the sequence contigs included into VIRGO, including reference data sets: i) complete VIRGO database, ii) 212 *in-house* sequenced vaginal metagenomes, iii) 53 HMP DACC vaginal metagenomes [32], iv) all HMP urogenital reference genomes, v) 277 genomes of bacteria isolated from vagina, reproductive or urinary system deposited in GenBank, and vi) 139 genomes of urogenital bacteria from HMP DACC database [15].

**Additional file 2: Table S4.** Table showing counts of the non-redundant genes in VIRGO by taxonomic groups in both species and genera.

**Additional file 2: Table S5.** The vaginal fungal database that includes 5 vaginal yeast species in 40 genomes and the abundance of detected fungal and phage in metagenome samples.

**Additional file 2: Table S6**. Annotation and alignment of a Jaccard orthologous clusters (JOCs) involved in vaginolysin. This JOCs was in one protein family in VOG that contains multiple genes, annotation information is in **A**. Multiple sequence alignment of this protein family was performed in T-Coffee [95], was used to access the alignment quality (**B**).

**Additional file 2: Table S7.** Examples of cell surface-associated proteins of *L. iners*. Two Jaccard orthologous clusters (JOCs) involved in this function were retrieved from VIRGO. (**A**) the JOC was recognized to have LPXTG motif; (**B**) the JOC that harbor motif YSIRK.

**Additional file 2: Table S8.** The statistics of number of non-redundant genes identified in a metagenome and the depth sequencing for samples in different CSTs.

**Additional file 2: Table S9**. Examples of tryptophan production-related gene cataloging using VIRGO. It includes three essentials genes Tryptophanase (*TnaA*), Tryptophan synthase beta chain (*TrpB*), and Tryptophanyl-tRNA synthetase (*TrpS*) in gene name, gene richness, gene annotation, to demonstrate the profiling of a specific function of interest and its taxonomic distribution.

**Additional file 2: Table S10**. Summary of the 7 vaginal bacterial species with gene content characterized using VIRGO to determine the diversity of individual populations. It includes four *Lactobacillus* species (*L. crispatus, L. iners, L. jensenii*, and *L. gasseri)*, as well as three additional species common to the vagina (*G. vaginalis, A. vaginae* and *P. timonensis*). Reads mapping was performed using 1,507 *in-house* and publicly available vaginal metagenomes to VIRGO. Metagenomes that contained at least 80% of their average genome’s number of coding genes were included. Abbr: Av: *A. vaginae*; Gv: *G. vaginalis*; Pt: *P. timonensis*; Lc: *L. crispatus*; Li: *L. iners*; Lj: *L. jensenii*; Lg: *L. gasseri*.

**Additional file 2: Table S11.** List of accession numbers for genomes of the four *Lactobacillus* species including *L. crispatus, L. iners, L. jensenii*, and *L. gasseri* and three species including *G. vaginalis, A. vaginae* and *P. timonensis* used in intraspecies analyses.

**Additional file 2: Table S12.** Taxonomic distribution of pullulanase domain-containing proteins included in VIRGO.

### Availability of data and material

All database data and code were made freely assessable on https://github.com/Ravel-Laboratory/VIRGO. It includes Jaccard index clustering code, VIRGO non-redundant nucleotide gene database, VOG amino acid protein family database, curated taxonomy and functions information, and tutorials. Metagenomes used in the analyses are deposited at EBA ### (The list of accession numbers for the 1,507 vaginal metagenomes used in intraspecies analyses will be available upon acceptance of the manuscript).

### Authors’ contributions

B.M., J.R. designed the research. B.M., M.F., J.H., and J.R. performed the research. B.M., M.H. generated the data. B.M., M.F., J.H., and J.C. analyzed the data. B.M., M.F., R.B., and J.R. interpreted the data and wrote the paper.

## Acknowledgements

The authors thank Drs. Douglas Kwon and Matthew Hayward for their helpful assistance in analyzing African women metagenomes. The authors thank Dr. Nan Qin and Qian Xu for their helpful assistance in analyzing Chinese women metagenomes. Research reported in this publication was supported by the National Institutes of Allergy and Infectious Diseases and Nursing Research of the National Institutes of Health under award numbers U19AI084044, R01NR015495 and R01AI116799, and the Bill & Melinda Gates Foundation award OPP1189217.

