## Supplemental_Figures for "VIRGO, a comprehensive non-redundant gene catalog, reveals extensive within community intraspecies diversity in the human vagina"

**Additional file 1: Figure S1**. Boxplot of the proportion of sequencing reads after removing human contaminates from the samples between different Community State Types (CSTs). CSTs were defined as previously according to the composition and structure of the microbial community [29]. Plotted are interquartile ranges (IQRs, boxes), medians (line in box), and mean (red diamond). Significance value was calculated using Wilcoxon rank sum test using *ggsignif* R package [99]. Star sign (*) denotes the level of significance.

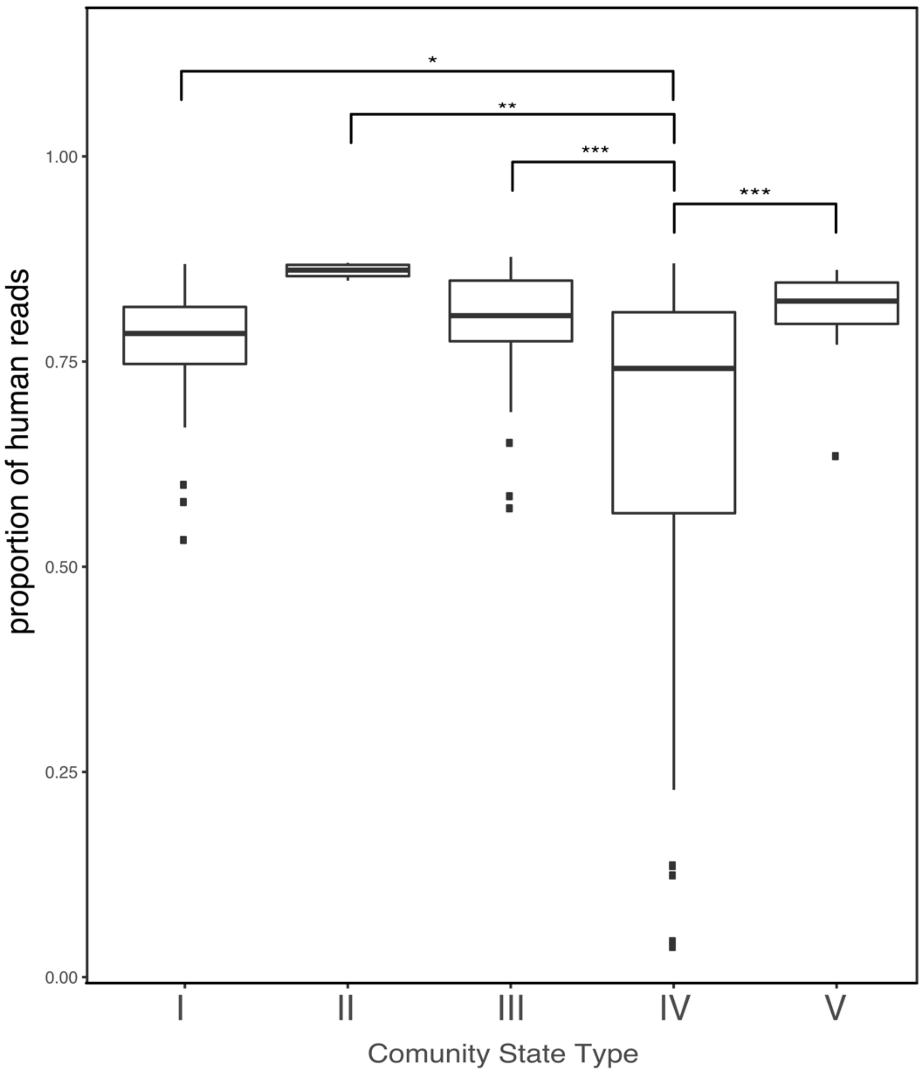

**Additional file 1: Figure S2.** Heatmap of relative abundance of the 50 most abundant phylotypes in the vaginal metagenomes used in this study. Ward linkage clustering is used to clusters samples based on their Jensen-Shannon distance calculated in the *vegan* package in R [100] according to the previous naming convention [29]. The sidebars indicate CSTs and gene richness category, respectively. Gene richness categories include high gene count (HGC) and low gene count (LGC), defined using the threshold of 10,000 genes per sample.

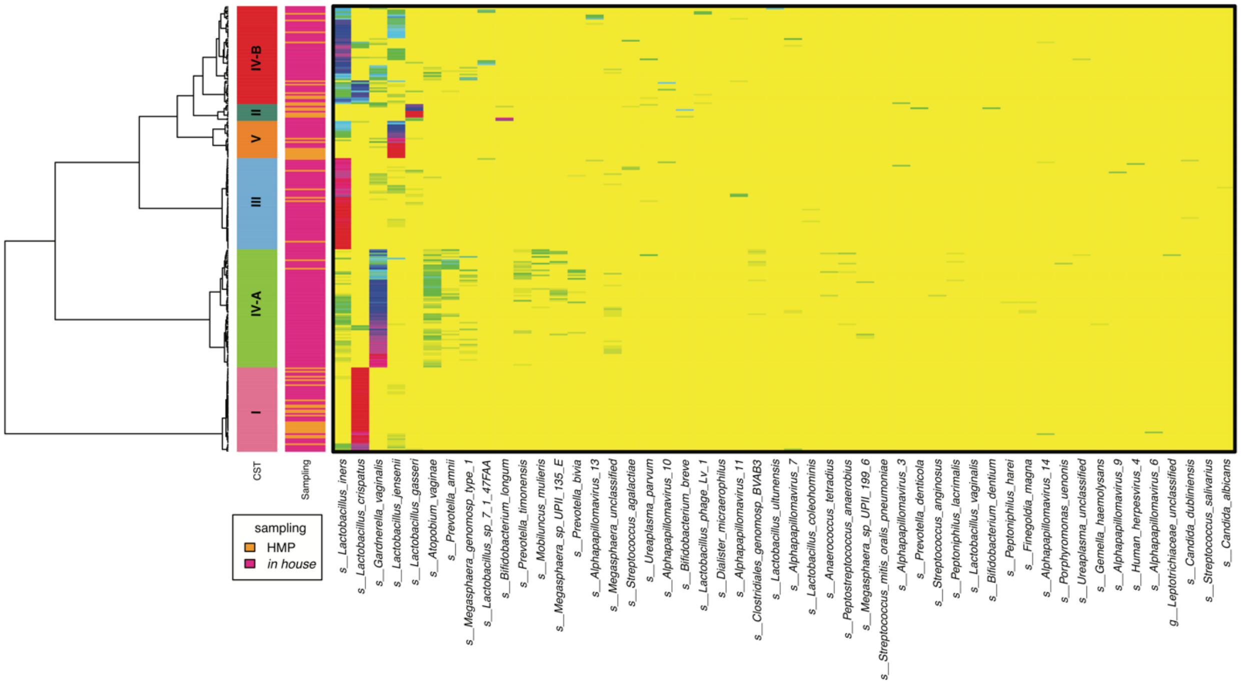

**Additional file 1: Figure S3**. Vaginal community accumulation curves and diversity estimate. (A) Accumulative diversity estimates with respect to sample size, for rarefied and extrapolated estimates using all samples; (B) accumulative diversity estimates with respect to sample size, for rarefied and extrapolated estimates using samples of different CSTs; (C) diversity estimate with respect to sample coverage, for rarefied and extrapolated estimates using all samples; (D) diversity estimate with respect to sample coverage, for rarefied and extrapolated estimate using samples of different CSTs. Community diversity estimates were computed using R package *iNEXT* [71] and *vegan* [72]. Sampling curve was either rarefied to smaller sample sizes or extrapolated to a larger sample size for species diversity estimate.

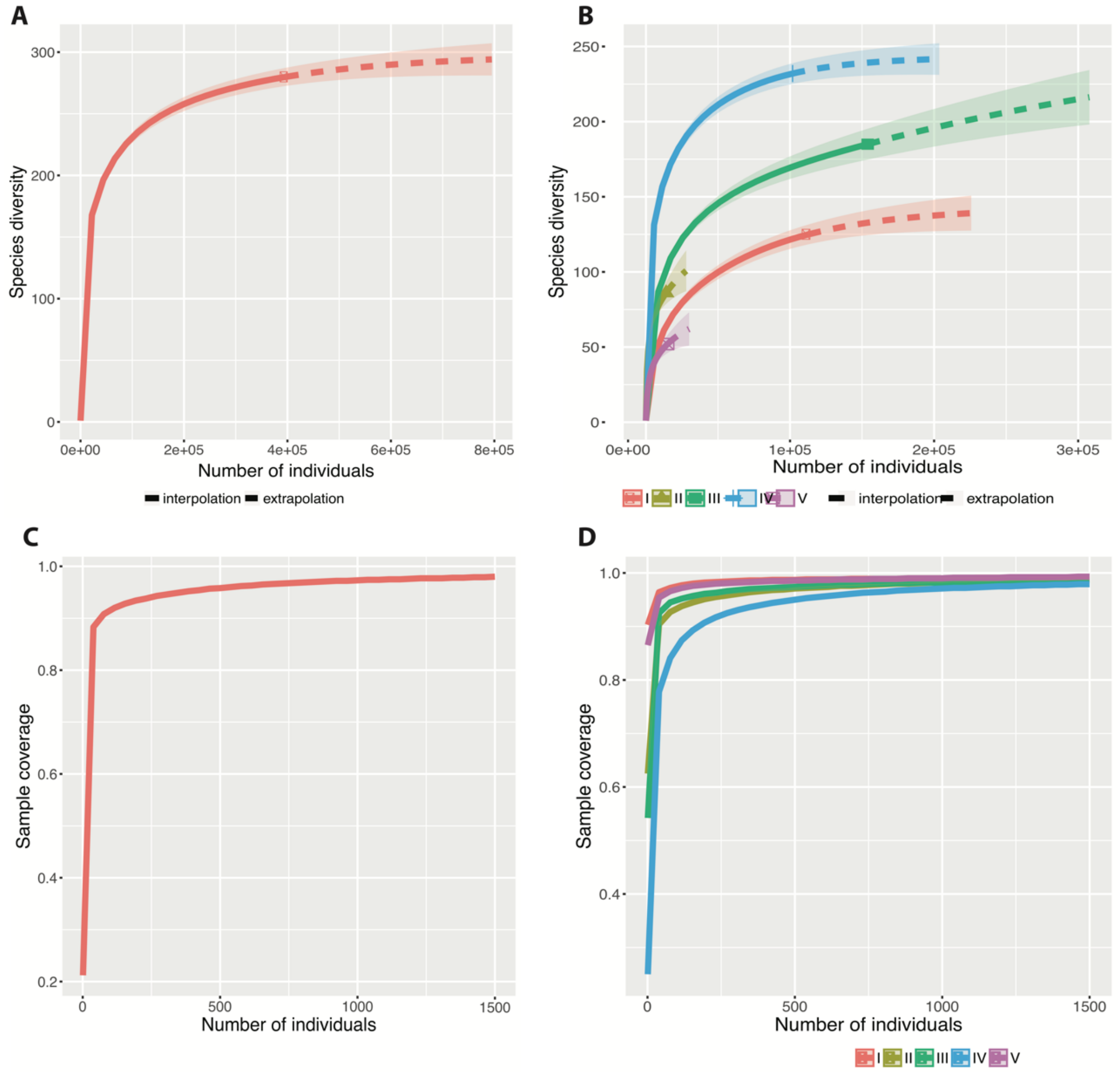

**Additional file 1: Figure S4**. Pie chart taxonomic distribution of reads that failed to map on VIRGO for vaginal metagenomes of African women from Gosmann *et al.* [30] in **A** and of Chinese women from [31] in **B**. The unmapped reads were compared to GenBank nt database [97] using BLASTN.

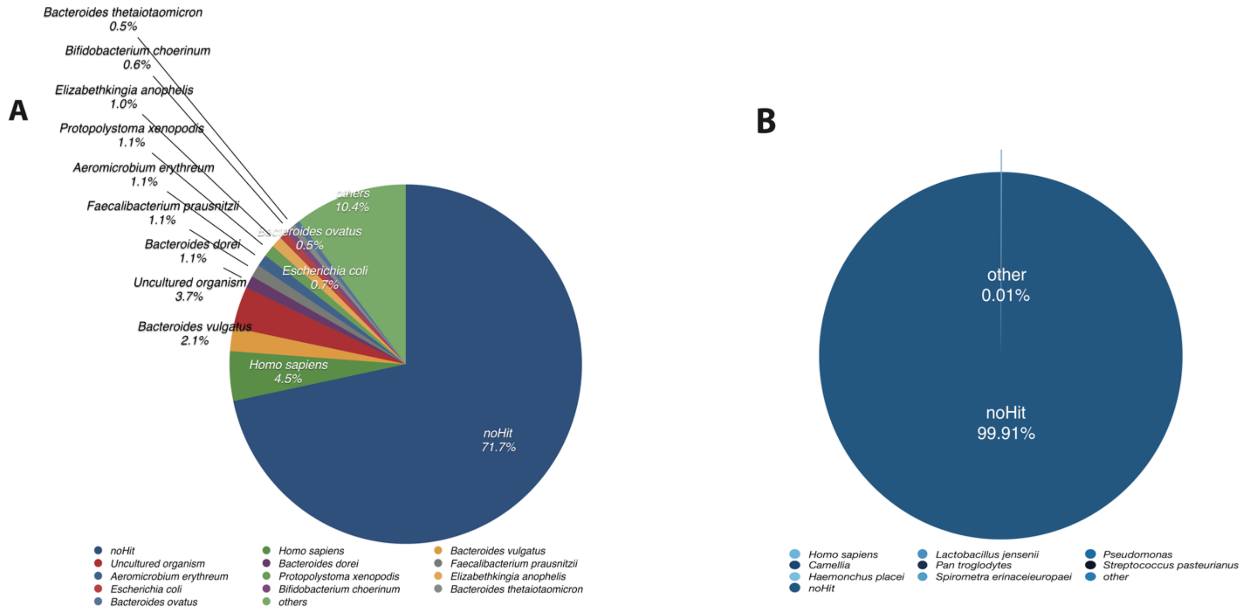

**Additional file 1: Figure S5**. Pipeline for data processing and integration for the construction of the human vaginal integrated non-redundant gene catalogue (VIRGO) and vaginal orthologous protein family groups (VOG). Metagenomes from 264 vaginal metagenomes and 416 genomes of urogenital isolates were processed, that including 212 *in-house* sequenced vaginal metagenomes. The procedures include pre-processing to remove human contaminates, quality assessment, metagenome assembly, gene calling, functional and taxonomic annotation, gene clustering based on nucleotide sequencing similarity to form VIRGO, and Jaccard index coefficiency clustering of amino acid sequences to form VOG.

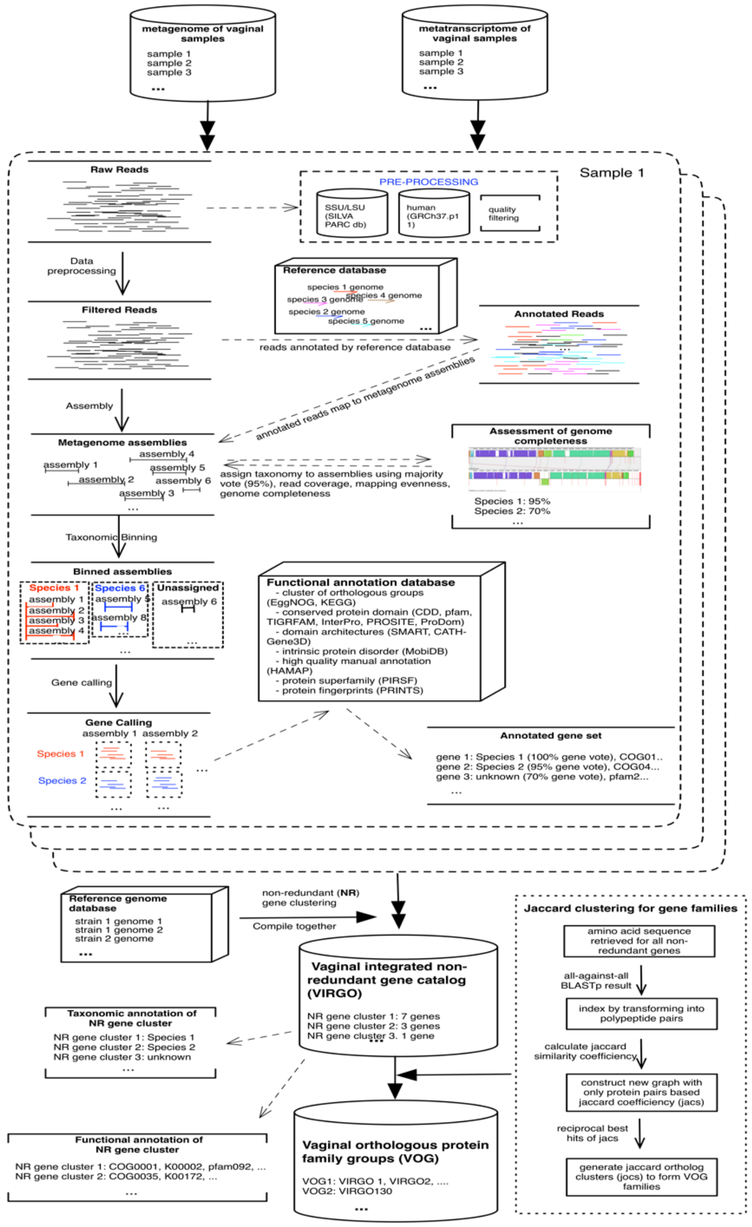

**Additional file 1: Figure S6**. Proportion of the assembly length assigned taxonomically from the samples (**A**) among different community state types (CSTs) and (**B**) between different gene richness category. CSTs were defined as previously according to the composition and structure of the microbial community [29]. Gene richness category includes high gene count (HGC) and low gene count (LGC), defined using the threshold of 10,000 genes per sample.

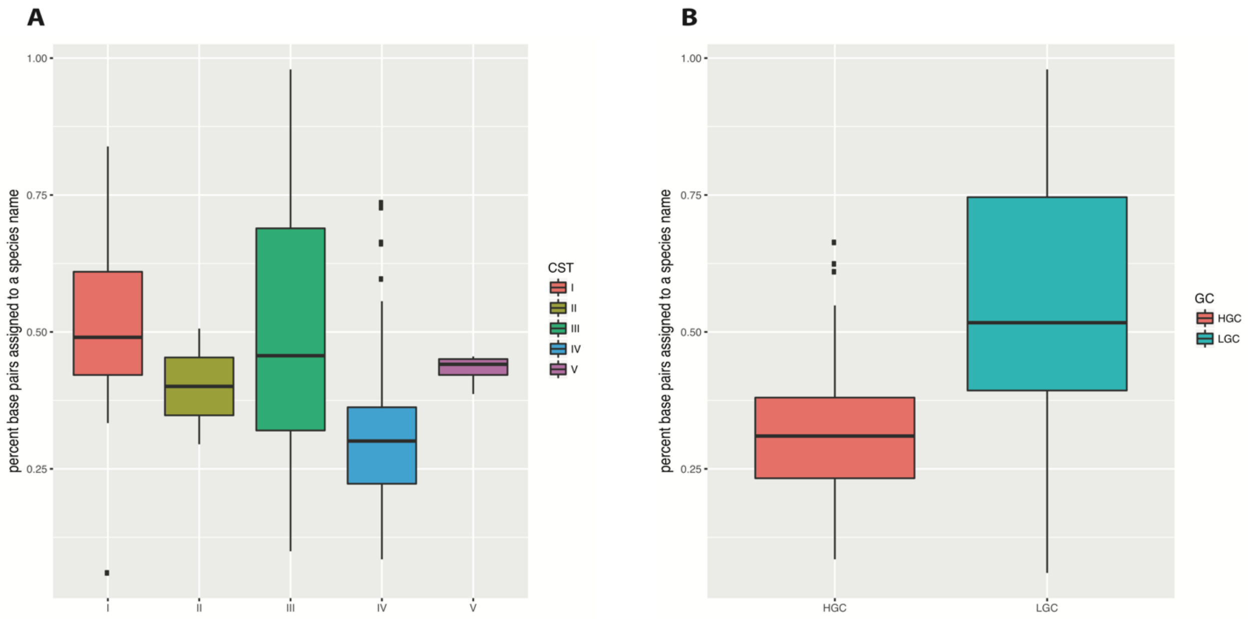

**Additional file 1: Figure S7**. Top 20 species with the most abundant gene content in VIRGO. The ratio of the gene content of a species over the entire community to the base 2. Plotted are interquartile ranges (IQRs, boxes), medians (line in box), and mean (red diamond).

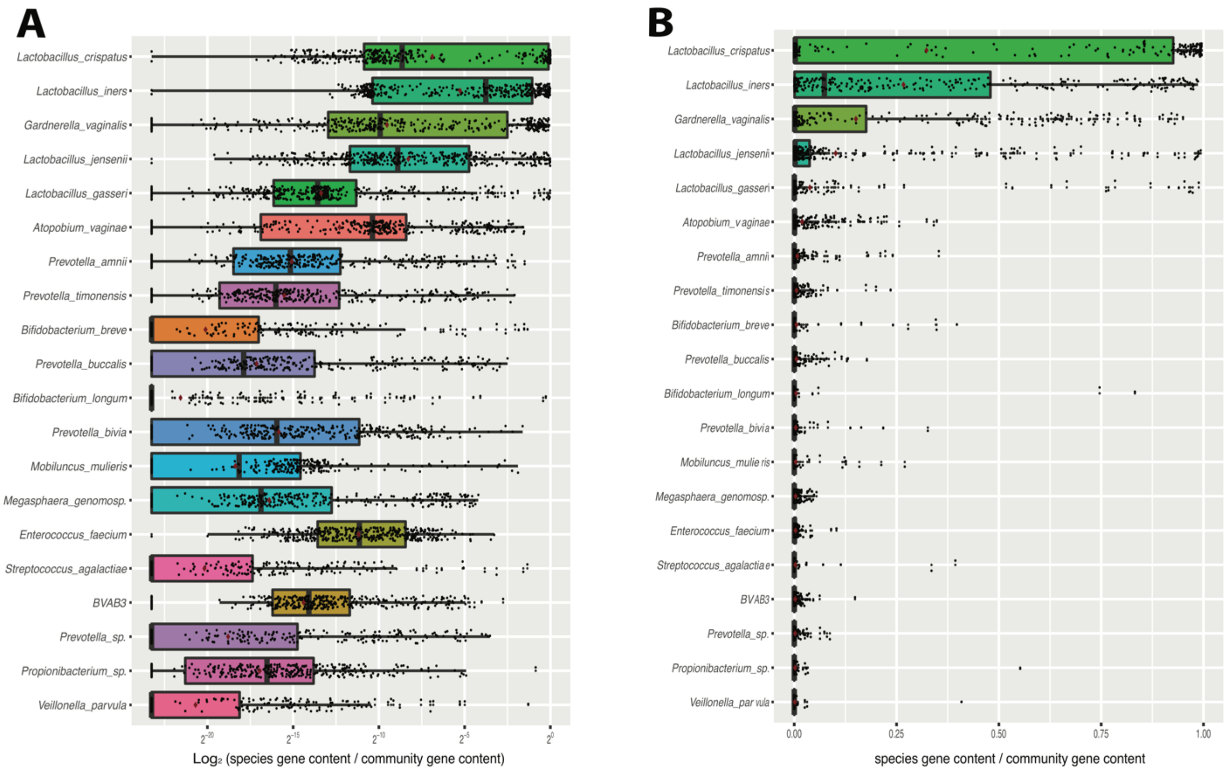

**Additional file 1: Figure S8**. Boxplot of the alignment scores of Jaccard orthologous clusters (JOCs) with multiple members. The alignment program T-Coffee [95] was used to access the alignment quality using alignment score.

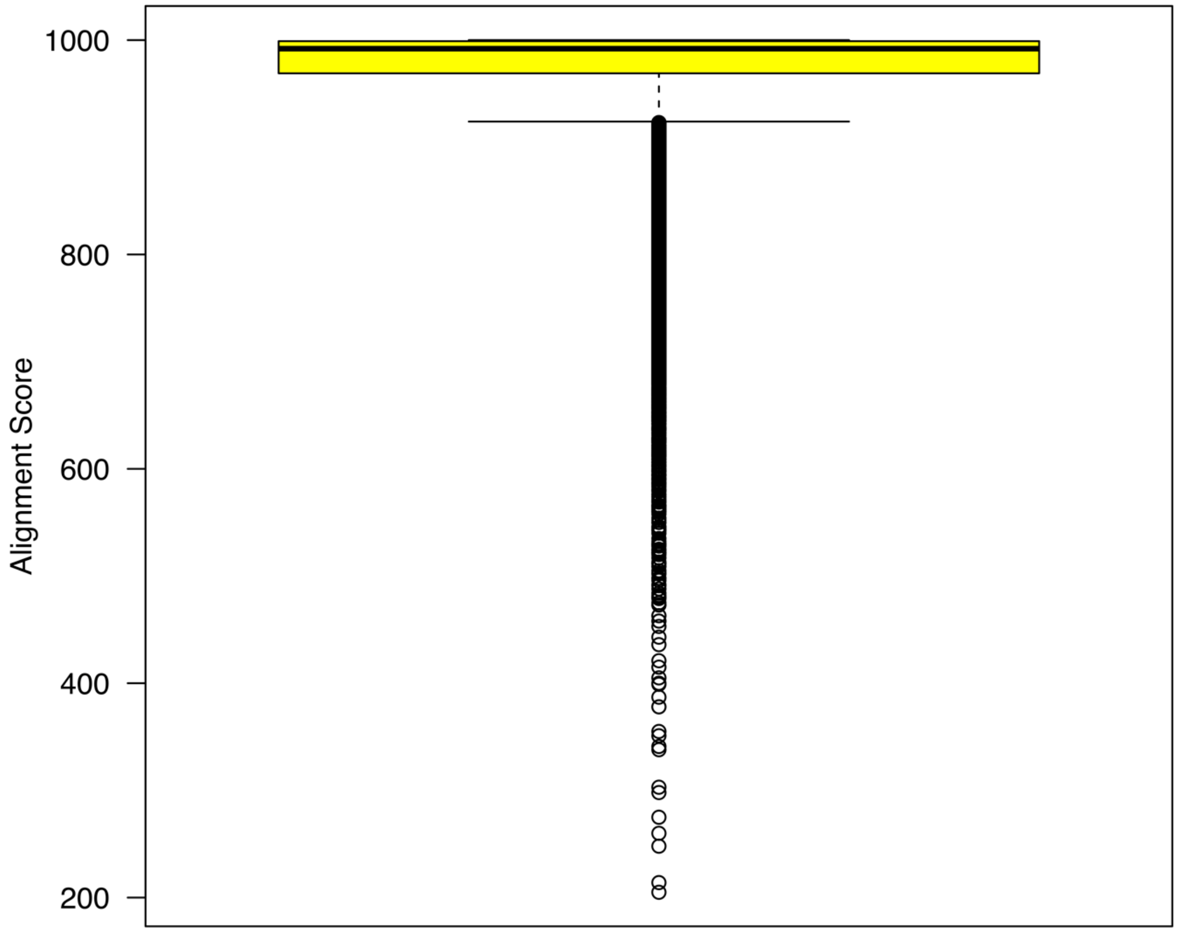

**Additional file 1: Figure S9**. Phylogeny that is demonstrative use of VOG to characterize the *G. vaginalis* cholesterol-dependent cytolysin (CDC) protein family. It shows the phylogeny of CDC-containing protein and alignment of domain 4 of the CDCs that is generally well conserved but contains a single divergent site, highlighted in yellow [38].

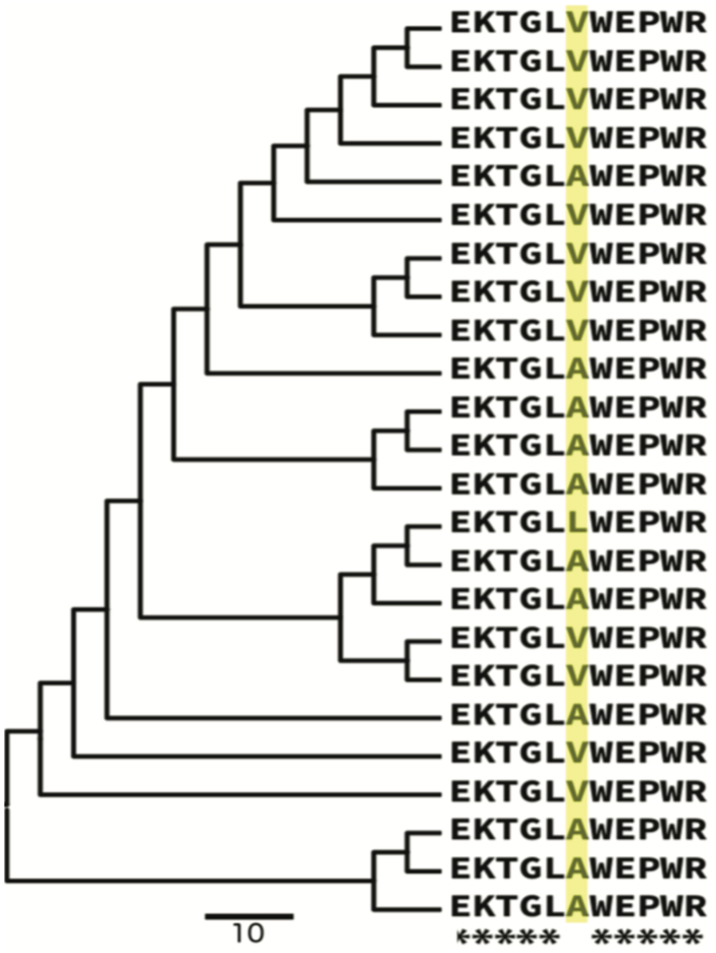

**Additional file 1: Figure S10**. Association plot of functional distribution of different gene count categories in vaginal microbiome. Functional category was defined using EggNOG (v4.5) [77] functional category. A Cohen-Friendly association plot [101, 102] was produced in statistical package *vcd* in R [103] to indicate deviations to indicate deviations from independence of CSTs and functional distribution. Mosaics display was shown, where the cells are shaded in proportion to standardized residuals, where the positive value (blue) is the observed frequency is substantially greater than would be found under independence, and the negative value (red) indicates cells which occur less often than under independence.

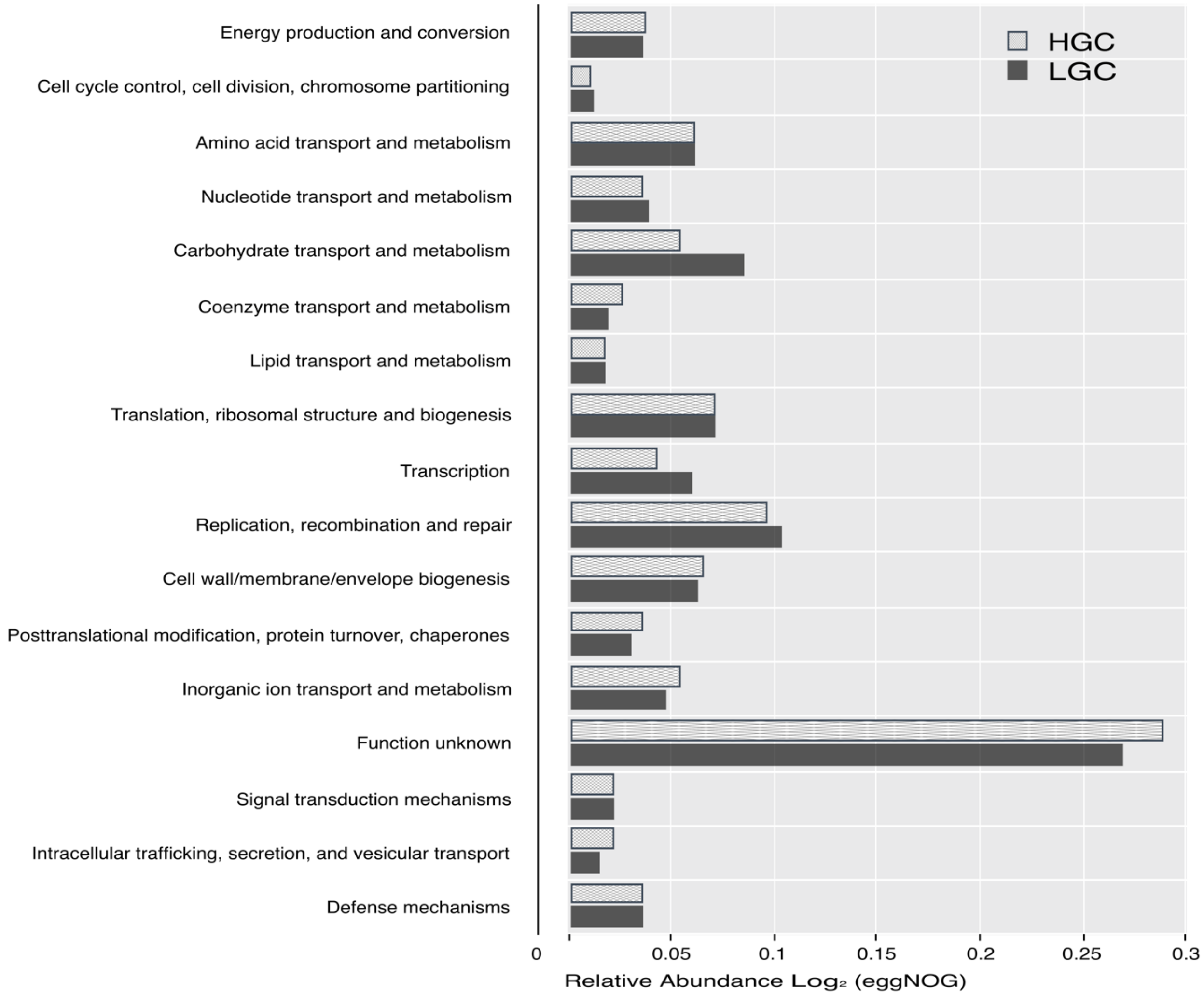

**Additional file 1: Figure S11**. Functional category of *L. iners* in different gene richness categories. Functional category was defined using EggNOG (v4.5) [77] functional category.

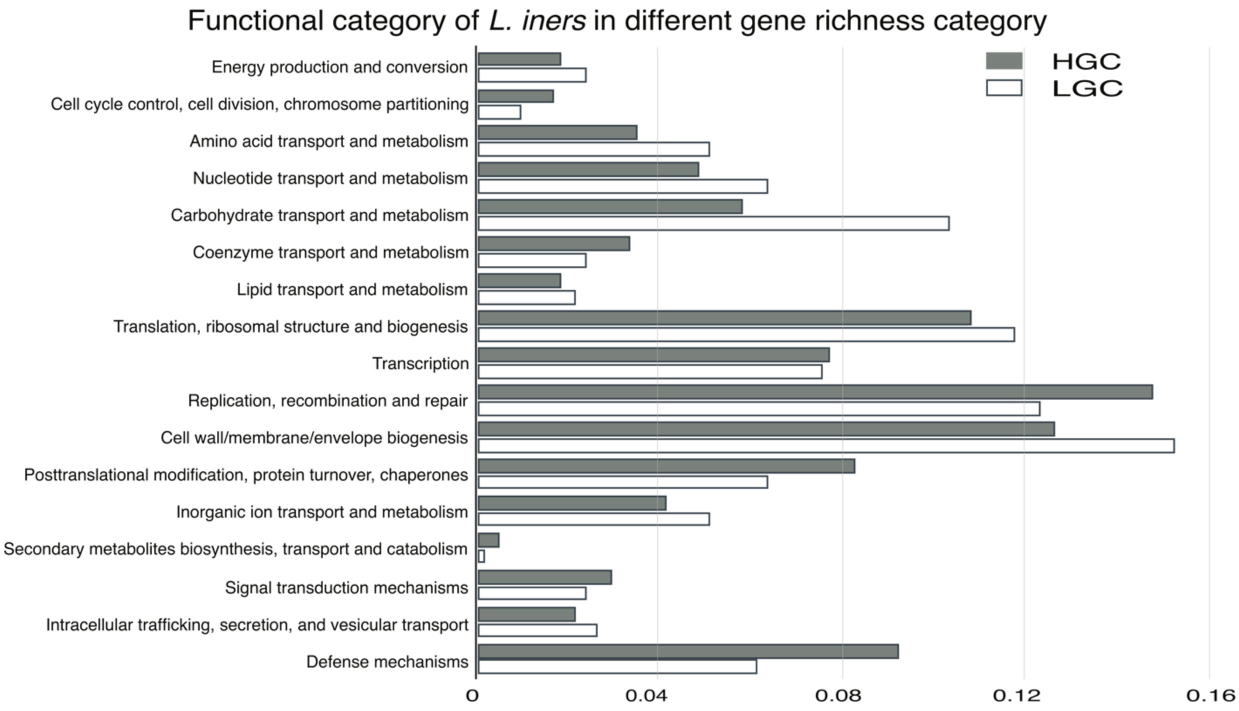

**Additional file 1: Figure S12**. Gene richness category and taxonomic distribution of tryptophan production-related genes in VIRGO. (**A**) Pie chart of the percentage of tryptophan production-related genes in different gene richness categories of HGC or LGC. (**B**) The top 10 most affiliated taxonomic groups of the tryptophan production-related genes.

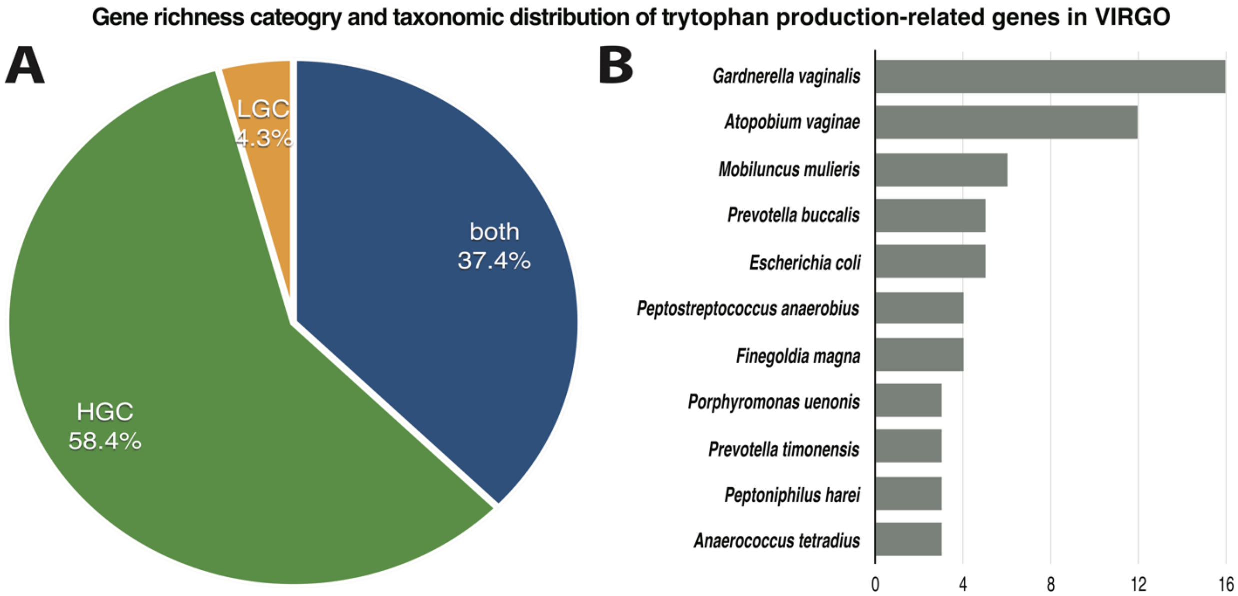

**Additional file 1: Figure S13**. Heatmap includes gene prevalence profiling of available genomes of vaginal isolates and VIRGO-characterized metagenomes for (**A**) *L. iners,* (**B**) *L. jensenii,* (**C**) *G. vaginalis*, (**D**) *A. vaginae* and (**E**) *P. timonensis*. Hierarchical clustering of the profiles was performed using ward linkage based on Jaccard similarity coefficient. CSTs were defined as previously according to the composition and structure of the microbial community [29].

**(A)** *L. iners*

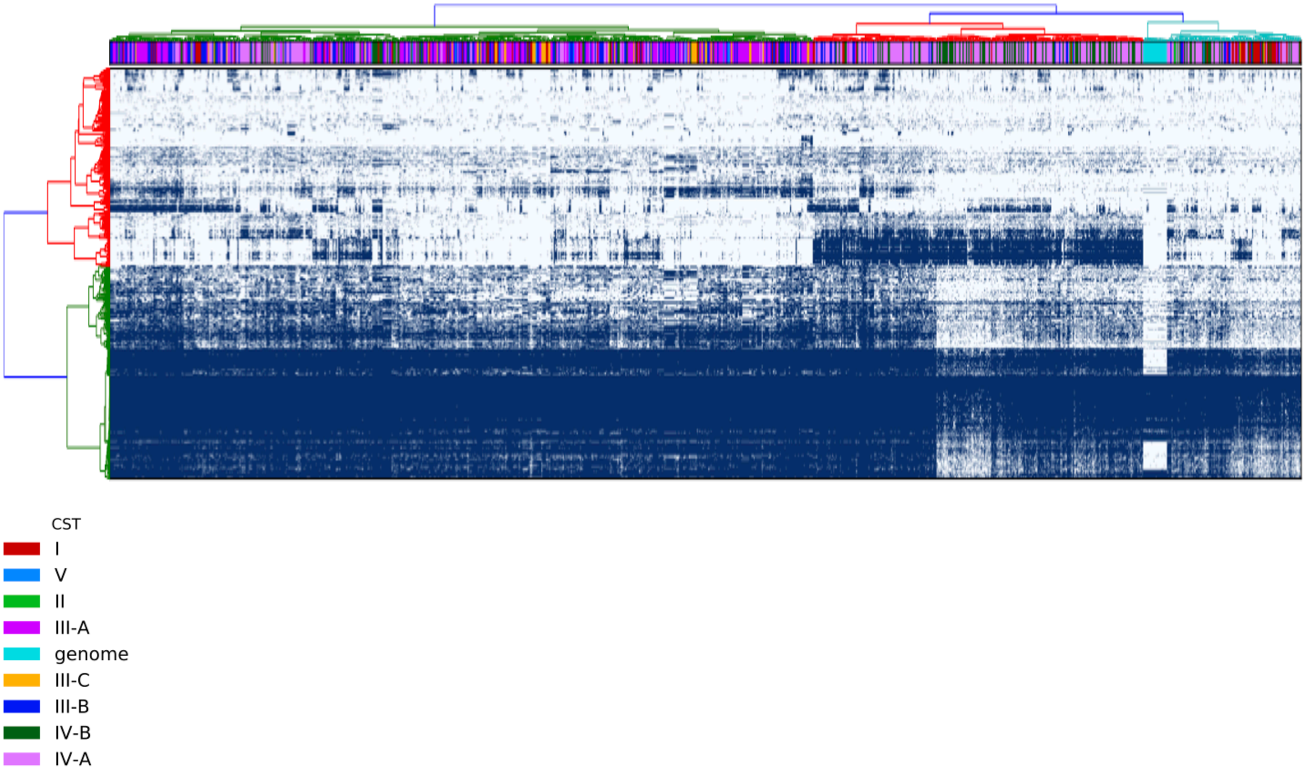

**(B)** *L. jensenii*

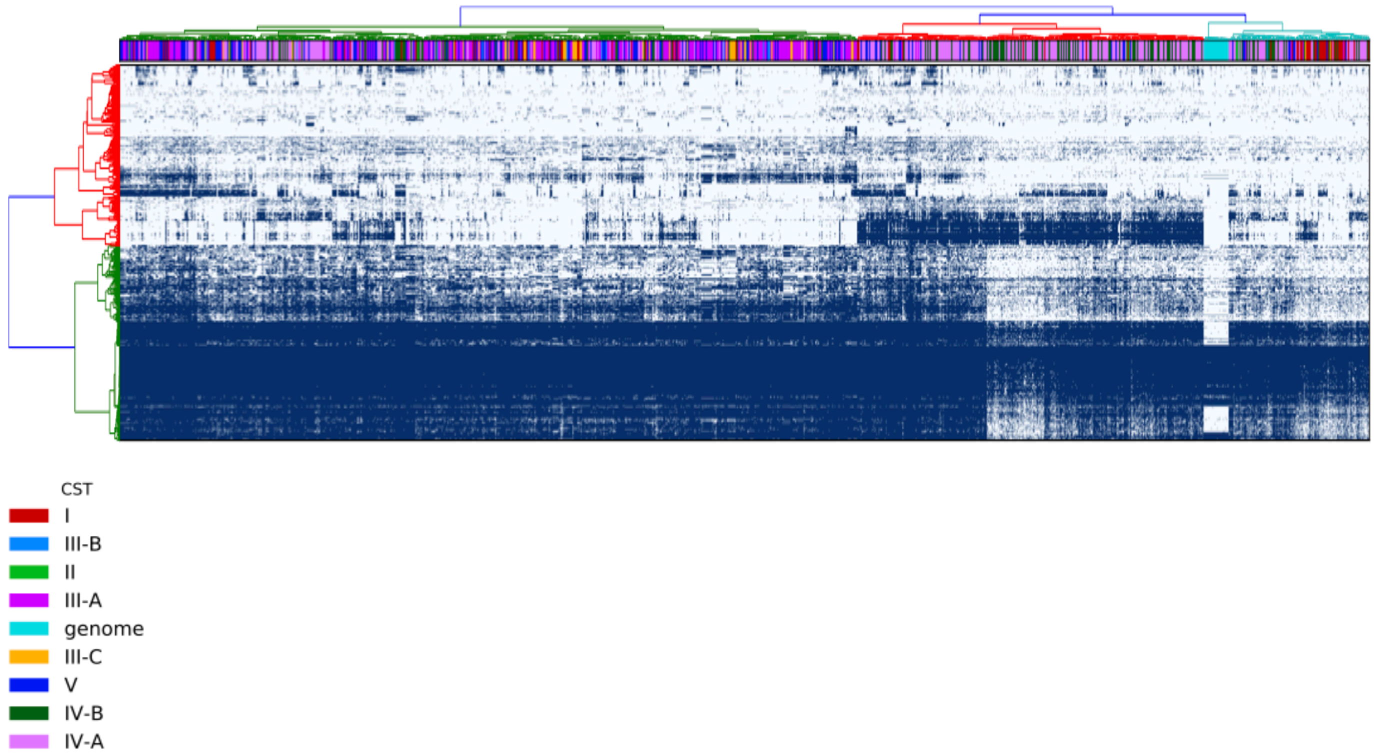

(**C**) *G. vaginalis*

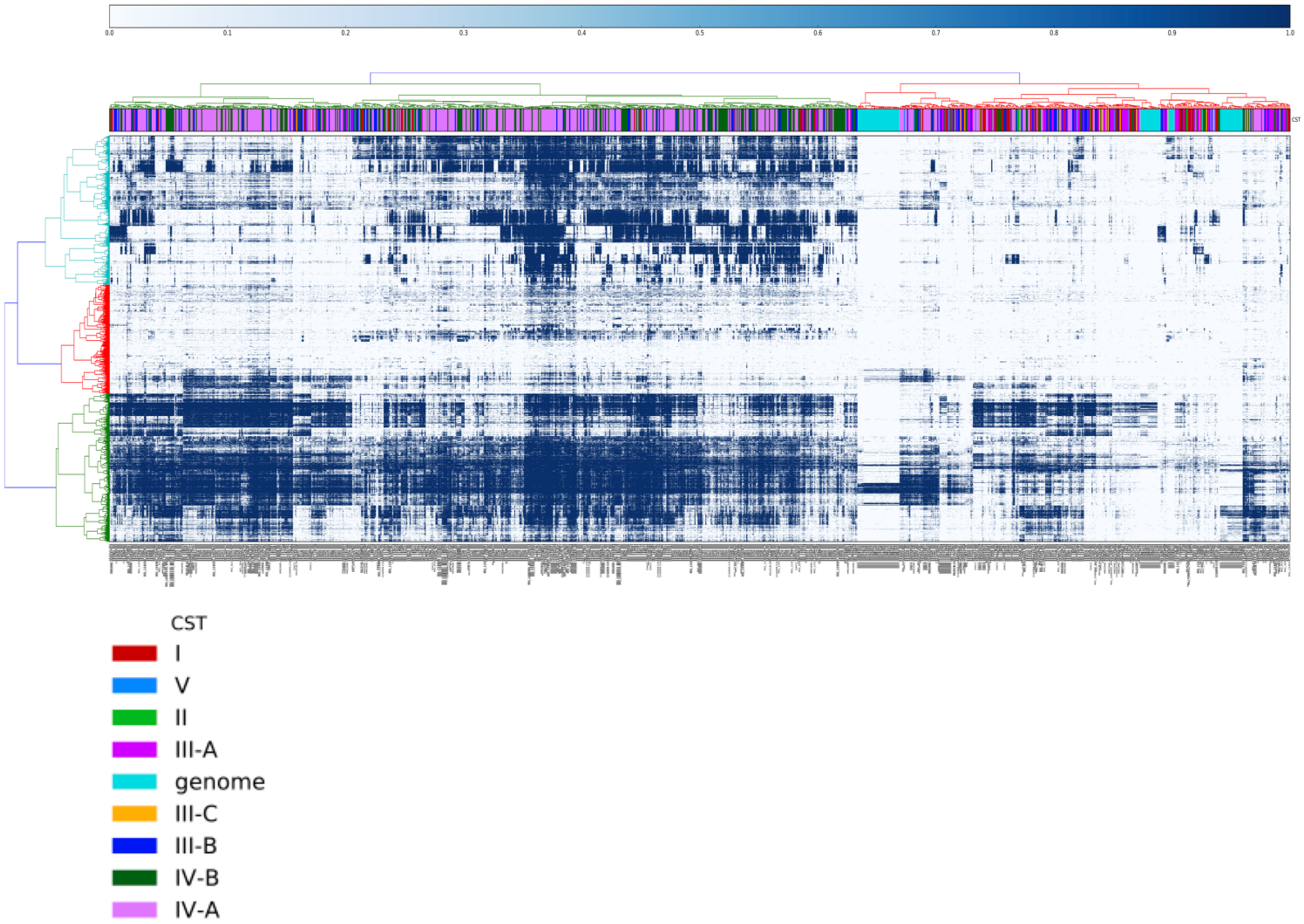

(**D**) *A. vaginae*

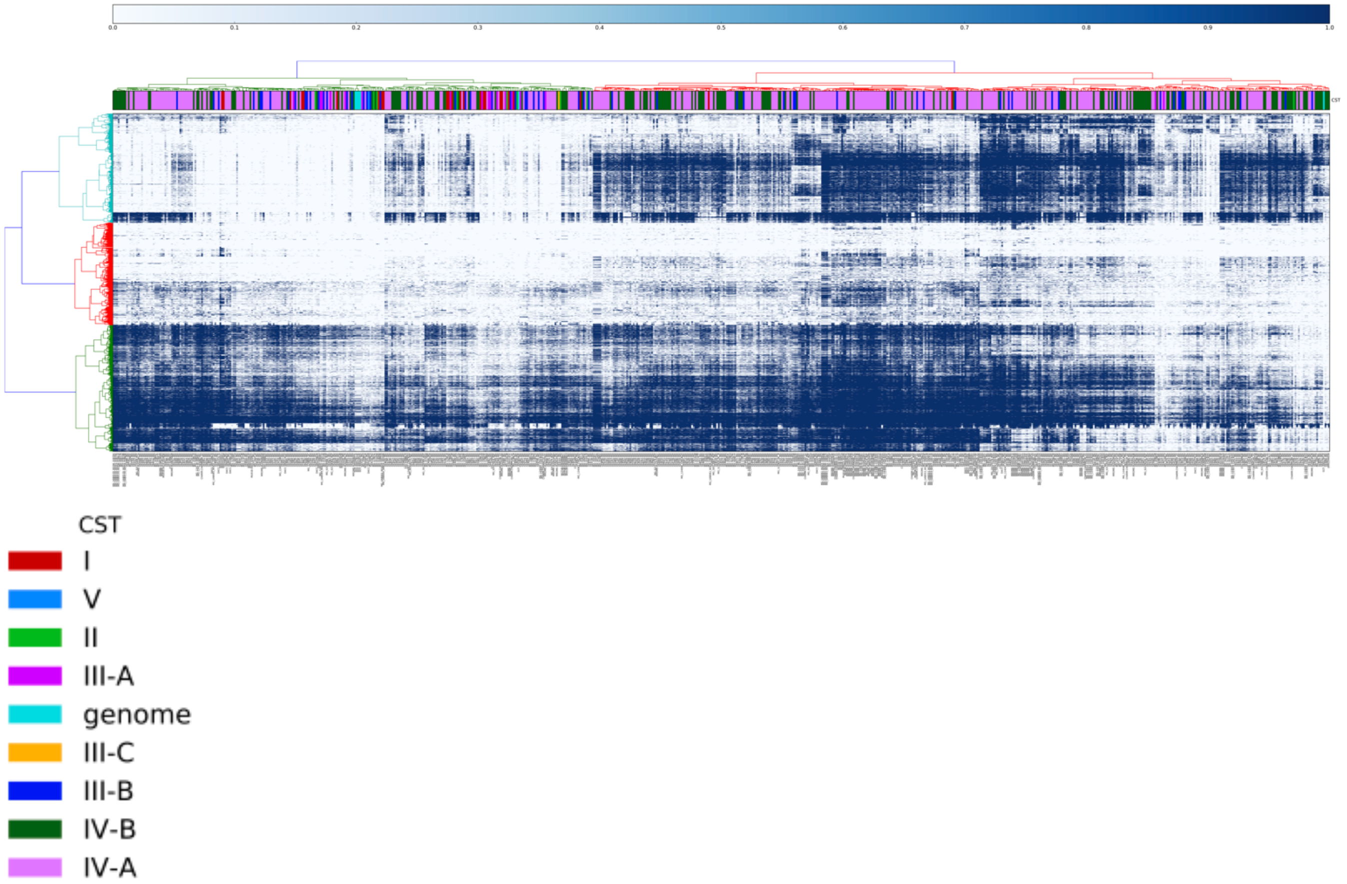

(**E**) *P. timonensis*

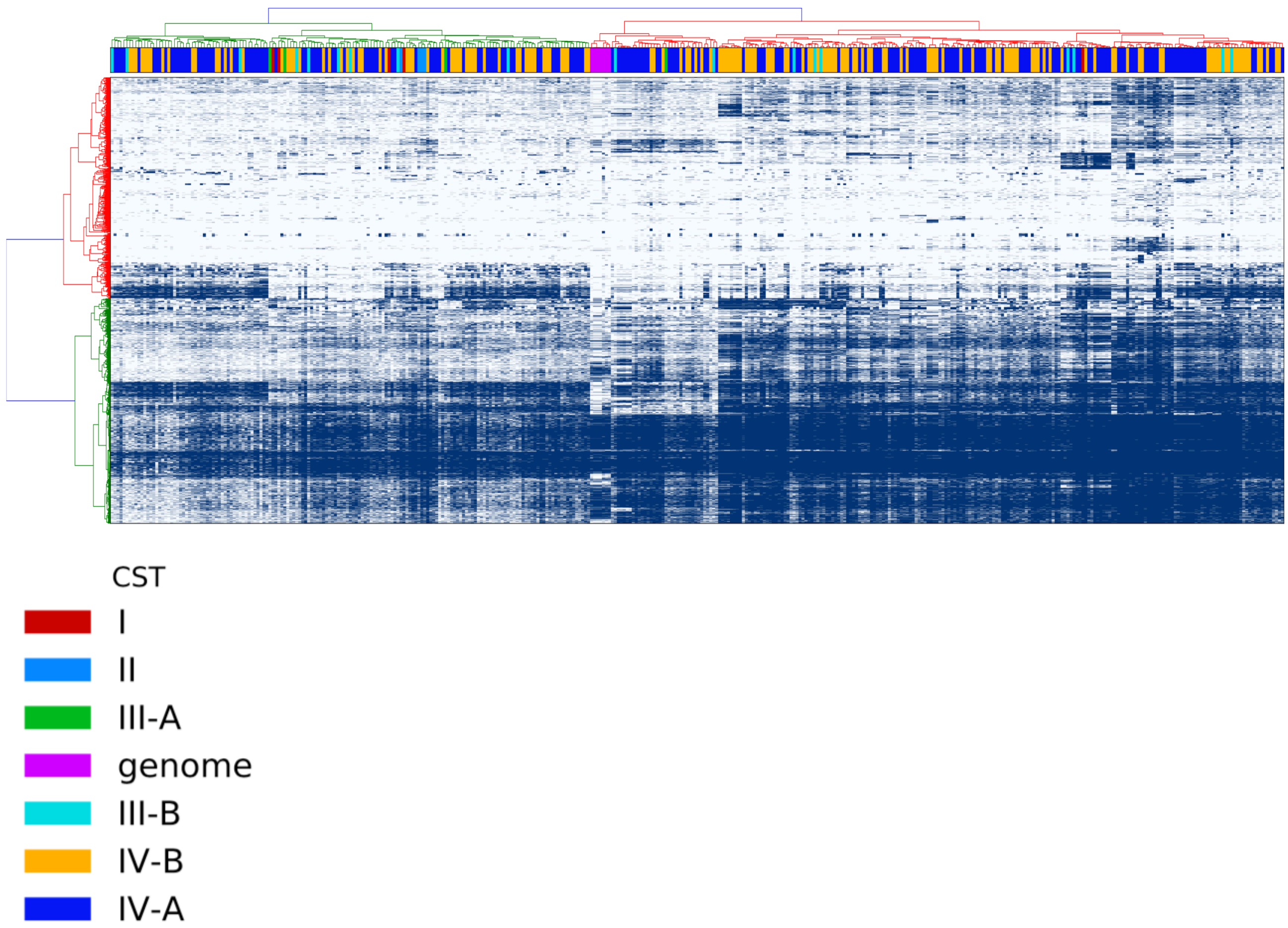
